# Laboratory rearing of the broad-nosed weevil *Scepticus tigrinus* (Coleoptera: Curculionidae) using artificial and plant-based diets

**DOI:** 10.1101/2025.06.24.660825

**Authors:** Yoshifumi Araki, Kei Matsubayashi, Toshiya Ando

## Abstract

*Scepticus tigrinus* Roelofs (Coleoptera: Curculionidae) and its related species are polyphagous weevil pests that damage various vegetables and crops. Despite their agricultural importance, scalable and straightforward rearing methods using artificial diets have not been established, limiting physiological and developmental studies necessary for effective pest control. The present study evaluated a commercially available artificial diet (F1675) for rearing *S. tigrinus*. We found that this artificial diet is suitable for the larval growth of *S. tigrinus*. However, the pupation rate was markedly low. To address this issue, we established a combined diet-rearing protocol in which larvae are fed with an artificial diet until the final instar and then exposed to raw plant pieces during the late final instar stage. This method significantly improved pupation success compared to the artificial diet alone (21.68%) and demonstrated higher pupation efficiency relative to labor input than the conventional raw plant-based rearing. Notably, final instar larvae pupated without feeding on the plant material, suggesting that non-nutritive cues—possibly chemical signals from the plants—may trigger pupation. The proposed combined diet protocol offers a practical approach for mass rearing and facilitates physiological and genetic studies of *S. tigrinus* and related weevil pests.

## Introduction

*Scepticus tigrinus* Roelofs (Coleoptera: Curculionidae) is a flightless, polyphagous beetle on sandy seashores facing the Sea of Japan (Yamashita et al., 2019). Broad-nosed weevils of the genus *Scepticus*, including this species, are known as polyphagous herbivores and pests of various vegetables and crops in Japan (Oida et al., 2021). Adults and larvae damage the plants’ above- and below-ground parts, respectively. In the wild, the life cycle of *S. tigrinus* is univoltine; overwintered adults become active in late April and lay eggs from early May to late July, and new adults emerge from late August to late October. It takes approximately 90-120 days from egg laying to emergence (Katsumata, 1934). For pest control research on *S. tigrinus* and some related species, stable rearing methods using living plants or raw plant pieces have already been established (Masaki & Sugimoto, 1991; Oida et al., 2018). However, the existing methods are labor-intensive and unsuitable for large-scale rearing when investigating the physiology and genetics of *Scepticus* weevils for effective pest control and other basic research. One of the most critical aspects of rearing is maintaining a clean environment to prevent mold contamination in the culture cases. To this end, raw plant-based diets require frequent replacement every few days. This frequent replacement demands substantial human effort, hindering the ability to conduct unrestricted and dynamic research.

The use of artificial diets with high preservability can solve some of the problems of the existing rearing method. In some weevils, rearing systems using artificial diets have been established, such as in the West Indian sweet potato weevil, *Euscepes postfasciatus*, and the olive weevil, *Pimelocerus perforatus* (Shimoji & Kohama, 1996; Yoshida et al., 2018). However, no practical rearing system with artificial diets has been reported for the genus *Scepticus*. Therefore, we focused on an artificial diet, F1675 (Frontier Agricultural Science, Newark, DE, US), which can be applied to a polyphagous weevil pest of North America, *Diaprepes abbreviatus* (Lapointe et al., 2008; Lapointe & Shapiro, 1999), which was initially developed for the blue-green citrus root weevil, *Pachnaeus litus*.

In the present study, we developed a novel rearing system using an artificial diet and silica sand that enables stable maintenance of *Septicus* weevils until the final instar, despite high mortality at the pupation stage. To address this limitation, we further established a combined rearing protocol in which larvae are transferred to a raw plant diet at the final instar, thereby significantly improving pupation success. This system provides a robust platform for large-scale laboratory rearing and will facilitate future studies on the developmental physiology and genetics of *Scepticus* weevils.

## Materials and methods

### Individual rearing system using artificial diets

We prepared an individual rearing system in a ϕ50 mm × 9 mm plastic Petri dish (Falcon 1006, Corning, NY, USA) with 6 mL of the artificial diets processed as follows: we combined 250 mL of water with 2.03 g of Agarose S (Nippon Gene, Tokyo, Japan) and 59.375 g of commercially available diet for insects, and autoclaved mixture at 105°C for 10 min. Then, 0.5625 g each of methyl-paraben and benzoic acid, which were dissolved in 99% ethanol, were added to the diet mixture as a preservative. After dispensing the artificial diet mixture, they were solidified and dried for 4 hours at room temperature and stored at 4°C until use.

### Adult weevil husbandry

Adult *S. tigrinus* were collected at a coastal area outside the special protection zone of Tottori Sand dunes (Tottori Prefecture, Chugoku district, Honshu, Japan) and maintained in our laboratory to establish a laboratory strain. Adult weevils from the wild were placed in pairs into plastic 1-ounce cups with silica sand No. 7 and were provided with fresh organic strawberry *Fragaria* × *ananassa* leaves and artificial diet pieces. Adults that emerged in the laboratory were kept in sealed plastic containers; the bottom of the container was lined with paper towels, and 1.5 cm squares of non-woven gauze were provided as an egg-laying substrate. Eggs were collected two times per week. Unless otherwise noted, insects at all life stages were kept under the following conditions: 25±1°C, 16L-8D photoperiod, and 60-70% RH.

### Larval rearing setup using the artificial diet in 2 mL microtubes

We collected eggs from the adults of wild *S. tigrinus* between October 2nd and December 28th, 2023, and gave them the young root of broad bean until their body length reached 3 mm (about 3-4 weeks). The surviving larvae were transferred to 2 mL microtubes filled with 750 μL of artificial diet on the bottom. 1,250 μL of silica sand No.7 was put on the larva (Figure 1a). The microtubes with artificial diets were replaced every 2 weeks or when they were dried up. They were maintained at 25±1°C, in total darkness, and at moderate humidity (the actual value was unknown) with saturated salt solutions. Emerging adults of this generation formed the first laboratory generation and served as parents in subsequent experiments. We used the *lm* function in the R software (version 4.4.2; R Core Team, 2024) to estimate the regression line and assess the correlation between the number of introduced eggs and obtained larvae per cup.

**Figure 1.**
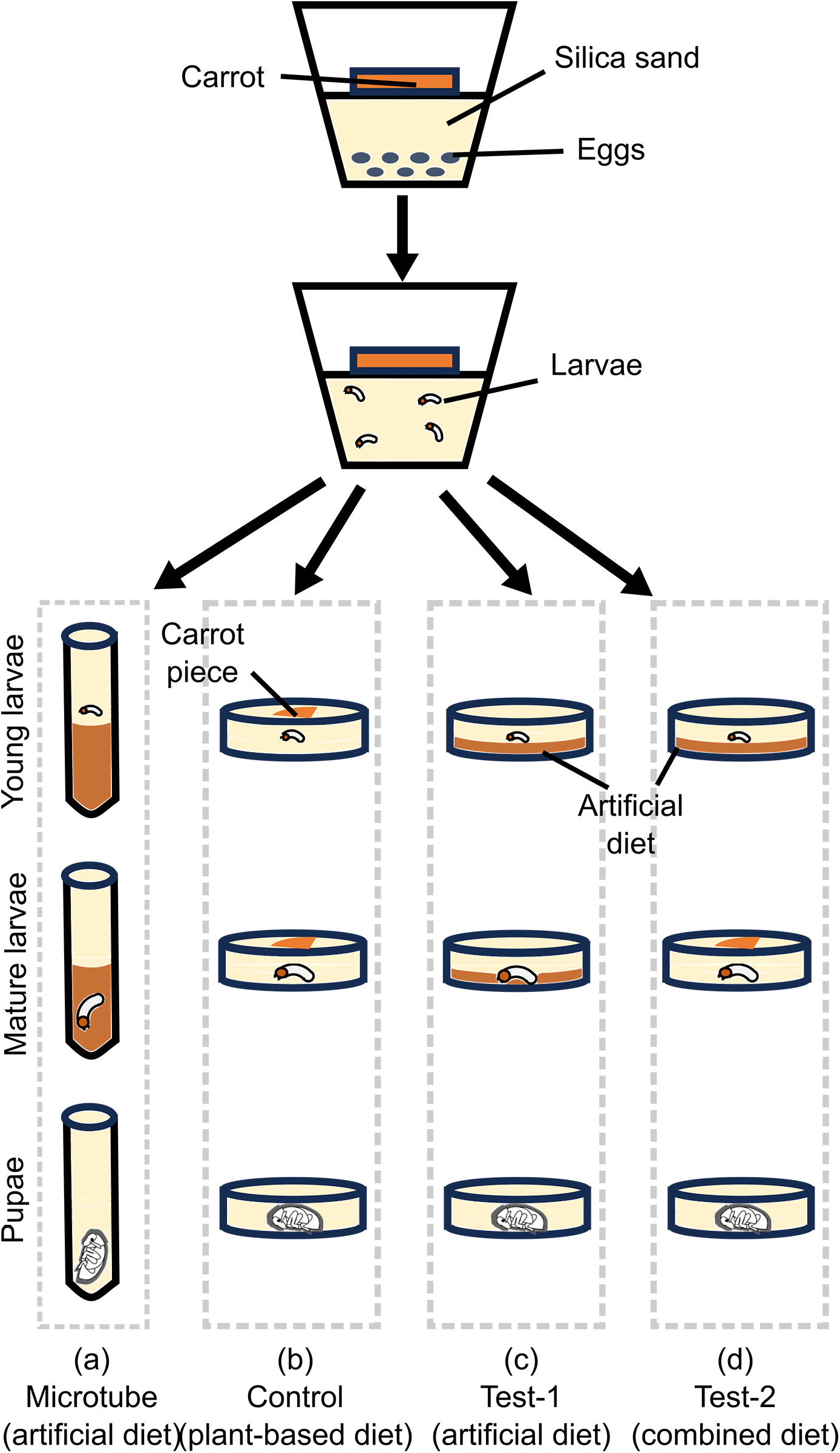
The workflow of the laboratory rearing of *S. tigrinus*. (a) Microtube rearing; (b) the plant-based rearing provided with carrot pieces only; (c) artificial diet rearing; (d) combined diet rearing provided with the artificial diet for the first 28 days and carrot pieces thereafter.

### Rearing setup for eggs, larvae, and pupae

Eggs were collected from adults in the first or second generation succeeded in the laboratory between July and December 2024. Eggs were collected in plastic 1-ounce cups and buried in silica sand and carrot *Daucus carota* subsp. *sativus* root pieces (carrot pieces) (Oida et al., 2021; Figure 1). The maximum number of eggs per cup was limited to 20. In 10 to 18 days after egg collection, the hatched larvae were divided into three groups to test different individual rearing conditions: the control group, the existing method using raw carrot pieces only; the test group, using the artificial diet only; the combination method group, using the artificial diet for the first 22-34 days and then carrot pieces (Figure 1).

In the control group, we followed the previous method using only carrot pieces (Figure 1b; Oida et al., 2021). Larvae collected before November 15th, 2023, were individually kept in Petri dishes with approximately 9 g of silica sand and a carrot piece. The carrot pieces were replaced every 3-4 days until pupation to prevent rot or mold. The pupae were transferred into new Petri dishes immediately after pupation, and enough sand and a carrot piece were used to maintain humidity.

In test group 1, we set up a rearing system using only the artificial diet (Figure 1c). We collected larvae on August 27th, 2024, and transferred them into the individual rearing system described above. Larvae and artificial diets were covered entirely with silica sand. In our initial trials, we noticed that the larvae could not be maintained without silica sand, as they sink and drown in the artificial diet. In addition, young weevil larvae could not be maintained on the completely flat artificial diet in silica sand. To help feed, we made a depression in the center of the artificial diet and inserted a larva into it (Figure 1). Diets were replaced every 28 days or when the diet became extremely dry.

In test group 2, we set up the rearing system combining artificial and raw plant diets (Figure 1d). Larvae collected between November 23rd and December 29th, 2024, were transferred first into the artificial diet during the first 22-34 days until the final larval instar and next into the carrot pieces. Managing the artificial and raw carrot diets was performed as described above. Pupae were transferred into new Petri dishes filled with silica sand and supplemented with preservative-added 4% agarose gel to keep humidity and avoid frequent mold contamination on carrot pieces.

In each experimental setup, we individually recorded the egg collection date, isolation, start of rearing using carrot pieces, pupation, and emergence. To compare pupation efficiency among rearing methods, we calculated the effort-based pupation efficiency (EPE) by dividing the pupation rate by the diet exchange effort, which is defined as the average number of days per diet replacement. In addition, we used the *t-test* function in the R package to conduct a Welch’s *t*-test to determine whether there was a significant difference in the number of days required for pupation between the plant-based and combined diet methods.

## Results and Discussion

### Rearing *S. tigrinus* larvae using artificial diet

To test the efficacy of the artificial diet F1675 in rearing *S. tigrinus* larvae, we set up a rearing system using plastic microtubes supplemented with F1675 and silica sand (Figure 1a). Of the 145 eggs collected, 69 larvae survived in the initial group rearing and were transferred to the microtube rearing. In this setup, 82.6% (57/69) of the individuals survived for 2 weeks after the start of the microtube rearing and became the final instar larvae, demonstrating a suitable setup for larval growth.

In contrast, larval development was markedly suppressed after two weeks, with only five individuals undergoing pupation and a single adult emerging (Table S1). After six months, 65.2% of the larvae (45/69) died. At this time point, 13% of the larvae (9/69) were still alive but had stopped development at the final instar or the prepupal stage. These results implied that pupation was impaired due to certain factors, such as inadequate nutrition or the lack of environmental cues necessary to trigger pupation. Therefore, we tested the effect of supplementing fresh carrot pieces and providing sufficient space (in a 5-mm Petri dish) on the 19 individuals that had survived for six months. Remarkably, 17 individuals underwent successful pupation, and 14 emerged as adults, including some already in the non-feeding prepupal stage (Table S1). All emerged adults matured sexually and laid eggs normally. During this experiment, we noticed that some larvae pupated without eating carrot pieces, suggesting that the low pupation rate in microtube rearing is due to the lack of environmental stimuli in the artificial diets or spatial availability in microtubes. These results indicate that this artificial diet is suitable for supporting larval development, apart from its tendency to cause pupation failure.

During the above experiments, we also noticed a trend of decreased survival with increasing egg numbers in group rearing (Figure 2), although the correlation was not statistically significant (p = 0.33). Cannibalism, which we frequently observed among early-stage larvae, appeared to be the primary cause of mortality. Therefore, in subsequent experiments, we limited each rearing container to a maximum of 20 eggs and restricted the rearing period to two weeks. This herbivorous weevil may have evolved cannibalistic behavior as an adaptation to the resource-limited conditions of the beach environment.

**Figure 2.**
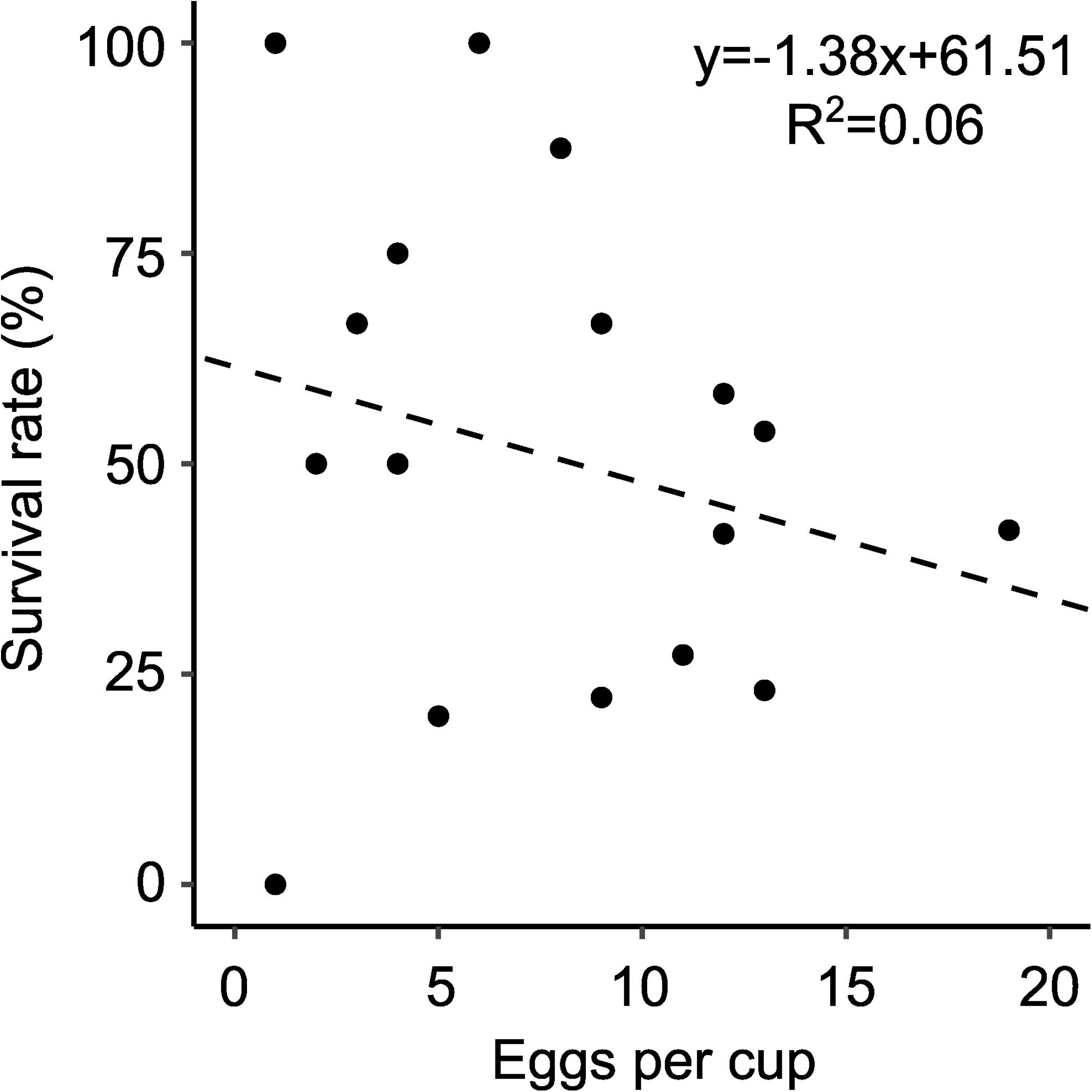
Correlation between the number of eggs per cup and survival rate. The dotted line indicates the regression line.

### Comparison of rearing methods with plant-based, artificial, and combined diets

We tested the efficacy of the above rearing protocol in which larvae are transferred to a raw plant diet at the final instar after artificial diet rearing. In this experiment, we collected a total of 599 larvae for comparative analysis and used 449 in the control group (plant-based diet), 7 in the test group (artificial diet), and 143 in the combination group (artificial and plant-based diets) (Table 1; Table S1). Of the individuals in the control (plant-based diet) group, 52.6% (236/449) pupated and 45.9% (206/449) successfully emerged. The mean duration from the start of individual rearing to pupation was 38.0 ± 9.3 days (95% CI: 36.8–39.2). In contrast, in the test (artificial diet) group, no individuals pupated and emerged as adults (0%, 0/7). Notably, of the individuals in the combination group (artificial and plant-based diets), 30.1% (44/143) survived in the final instar at the time of transfer to the carrot diet, 21.7% (31/143) pupated, and 6.3 % (9/143) emerged. The mean duration from the start of individual rearing to pupation was 50.0 ± 4.2 (95% CI: 38.3-41.7). Although the combined diet method resulted in lower pupation and emergence rates than the plant-based method, it proved to be a practical approach for laboratory rearing of *S. tigrinus* due to the reduced effort required for diet exchange.

**Table 1.** Numbers of total samples, larvae moved into carrot care, pupae, and emerged adults, periods requiring replacement carrot pieces (mean ± SD, 95% confident intervals), periods until pupation (mean ± SD, 95% confident intervals), and effort pupation efficiencies (pupation rate/periods requiring replacement carrot pieces).

|  | Larvae | In Carrots | Pupae | Adults | Period<br>with carrots | Period<br>to pupation | Effort-bas<br>ed<br>pupation<br>efficiency |
| --- | --- | --- | --- | --- | --- | --- | --- |
| Control | 449 | 449 | 236 | 206 | 37.99 ± 9.27<br>(36.78-39.20) | 37.99 ± 9.27<br>(36.78-39.20) | 1.38 |
| Test | 7 | - | 0 | 0 | - | - | 0 |
| Combined | 143 | 44 | 31 | 9 | 11.23 ± 3.44<br>(9.84-12.62) | 39.96 ± 4.18<br>(38.27-41.65) | 1.93 |

### Advantages and disadvantages of the combined diet protocol

The main advantage of the combined diet protocol lies in its reduced labor requirement. To quantify this, we calculated the effort-based pupation efficiency (EPE) score (as defined in the Materials & Methods section), which was higher in the combined-diet method (1.93) than in the plant-based method (1.44; Table 1).

On the other hand, in the above experiment, a decrease in post-pupation survival rate (29.0%, 9/31) was observed in the combination method, in contrast to a much higher rate in the plant-based diet method (87.3%, 206/236), which offset the superiority of the combined diet method in terms of EPE. This decrease in pupal survival rate seemed to be partly due to unexpected severe dryness, which we experienced during the experiment in the winter season because we needed to change agarose gel pieces for moisture maintenance more frequently. To test this possibility, we additionally measured the pupal survival rate in the same combined diet protocol from April 1st to June 2nd, 2025 (Table S1). In this experiment, we put the pupal-rearing container in a large plastic moist chamber to keep high humidity. The survival rate during pupa was 88.9% (16/18), comparable to that of the plant-based diet method (87.3%). Therefore, the superiority of the combination method is likely to remain valid even when the post-pupation survival rate is considered.

The disadvantage of the current combined diet protocol is still a lower pupation rate (21.68%). This lower pupation rate was due to the low larval survival rate (30.8%, 44/143), especially at the onset of the transfer into an artificial diet. During the survival check one month after diet transfer, we frequently observed that small larvae had died while remaining at the initial transfer site in the diet, while 70.45% (31/44) of the survived larvae successfully pupated (Table 1, Table S1). These observations imply that the primary cause of high mortality in the combining method is initial drowning due to excessive moisture leakage from the artificial diet to silica sand. The necessity of silica sand is the unique characteristic of the laboratory rearing of *Scepticus* weevils. To improve the survival rate at the initial larval transfer to an artificial diet, further improvement of the protocol or optimization of the composition of the artificial diet reported in other weevils would be necessary (Lapointe et al., 2010; Urasaki et al., 2009).

### Potential stimuli involved in pupation of *S. tigrinus*: an exploratory discussion

As mentioned in the first section, rearing with an artificial diet alone did not induce pupation in *S. tigrinus* despite supporting larval growth comparable to that under the plant-based diet (Welch’s t-test, t = –1.92, df = 58.88, p = 0.06; Figure 3). A notable finding in the present study is that the pupation of *S. tigrinus* was promoted when fresh carrot pieces were provided after rearing on the artificial diet. This finding implied that *S. tigrinus* requires environmental stimuli from the raw plant. A potential environmental stimulus for the pupation of *S. tigrinus* is chemical signals from plant roots. Root-emitted chemicals, such as volatile organic chemicals, CO_2_, and so on, mediate key aspects of insect larval behavior underground, including aggregation, attraction, and host plant choice (Arce et al., 2021; Johnson et al., 2016; Kojima, 2015; Nikoukar et al., 2025). A similar chemical signal might have been recruited to induce pupation in *S. tigrinus*. Future studies understanding the key signals to promote pupation in *S. tigrinus* will facilitate understanding of the physiology and life cycle of *Scepticus* weevils for pest control.

**Figure 3.**
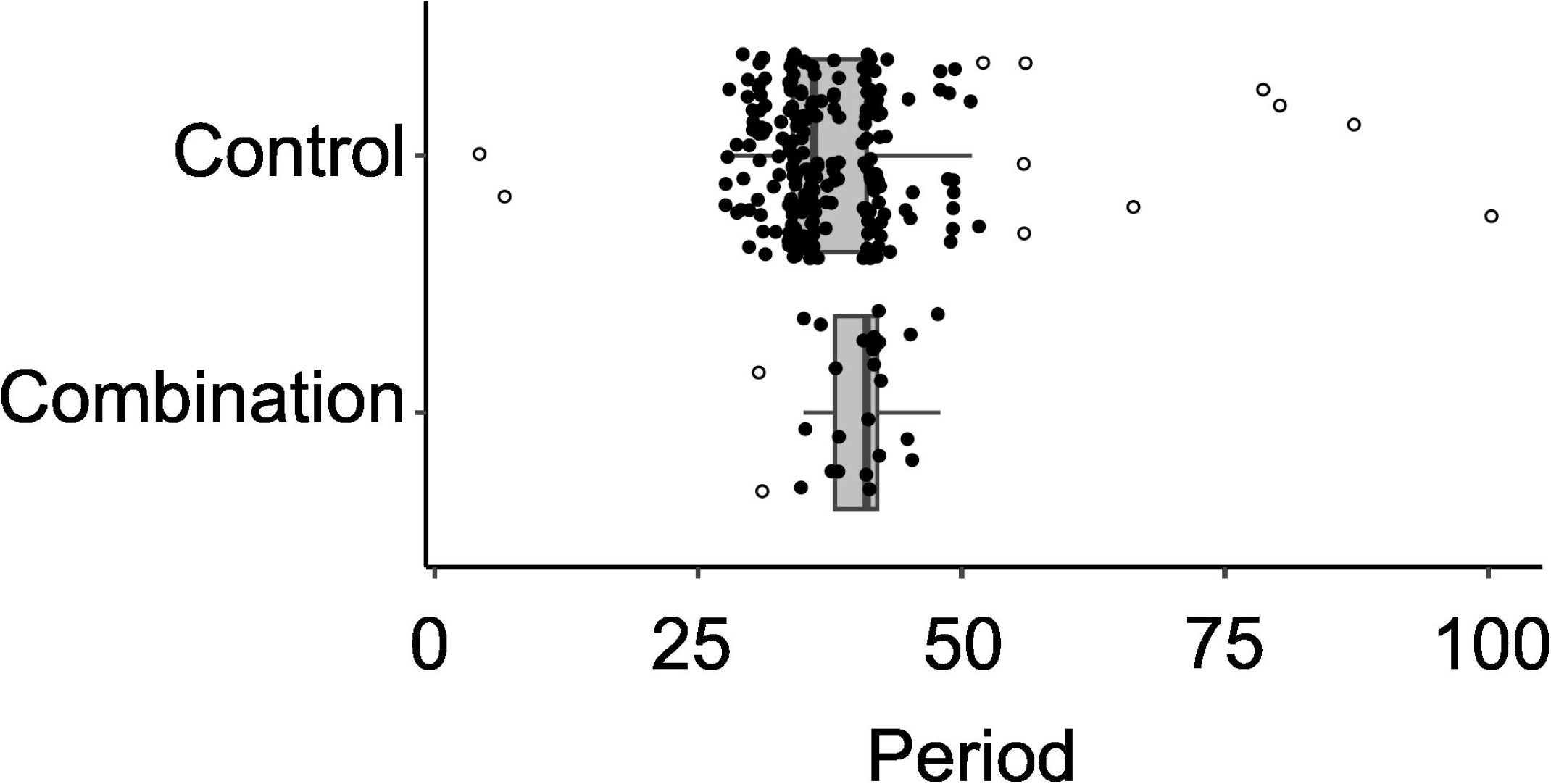
Box plot of the distribution of period to pupation; dot, individuals; circle, outliers; box, ranging from 25th to 75th percentile; vertical bar, median; horizontal bar, ranging from 0th to 100th percentile.

### Tips for the laboratory rearing of *S. tigrinus*

- A sufficient amount of silica sand is required to rear *S. tigrinus* larvae with an artificial diet. In the absence of sand, larvae become trapped in the wet diet and drown.
- Adult weevils can be maintained without silica sand, and eggs can be laid on paper and gauze. However, the number of eggs is increased when maintained with silica sand. Collect many eggs for rearing by sifting the sand with a stainless sieve with a 425 μm opening.
- Eggs and young larvae are highly delicate. When sifting silica sand to collect them, avoid shaking them directly on the stainless sieve by leaving a small amount of sand at the bottom. Use a wet, fine-tipped brush to gently pick up individuals from the sifted sand and transfer them to rearing containers.
- Larvae and pupae are sensitive to desiccation, while excessive humidity can lead to frequent contamination of mold and mites. Proper humidity control in the incubator, using saturated salt solutions or a compact dehumidifier, is essential for stable rearing.

## Conclusion

In the present study, we established a novel rearing protocol that combines artificial F1675 with a plant-based diet, enabling efficient rearing of the broad-nosed weevil *Scepticus tigrinus*. This protocol allowed the maintenance of a larger number of individuals while reducing the labor associated with the frequent replacement of the plant-based diet. We believe that this combined diet-rearing system will facilitate future studies on the ecology, development, and physiology of *S. tigrinus* and may contribute to developing pest control strategies based on rational design.

## Acknowledgments

We thank Dr. Hiroshi Oida at Hosei University for kindly providing us with helpful information on the plant-based diet-rearing method and Mr. Soichi Yeki for field sampling of *S. tigrinus*. We thank Mr. Takuma Sasaki, Mr. Takeru Toshito, Ms. Aya Fujikawa, and Ms. Yukiko Sado for their technical support. This study was supported by JSPS KAKENHI Grant Numbers 23H04309 and 23K19450 and JST CREST Grant Number JPMJCR24B1.

## Author contributions

Y.A., K.M., and T.A. conceived the study. Y.A. performed laboratory experiments and data analysis. Y.A. and T.A. wrote the manuscript with the input from K.M.

