## Supplementary material for "Laboratory rearing of the broad-nosed weevil *Scepticus tigrinus* (Coleoptera: Curculionidae) using artificial and plant-based diets": Table S1

Table S1. List of individuals and records of stage shifts. The dates were recorded in the YY/MM/DD format. Periods in the table indicate the death before that stage. Some adults that successfully emerged were sent to breeding.

| Individual_ID | Group | Egg Collection | Isolation | The Start of  Carrot Care | Pupation | Emergence | Death |
| --- | --- | --- | --- | --- | --- | --- | --- |
| TG1-5033 | Control | 2024/07/23 | 2024/08/06 | 2024/08/06 | 2024/09/09 | 2024/09/25 | Breeding |
| TG1-5034 | Control | 2024/07/23 | 2024/08/06 | 2024/08/06 | . | . | 2024/08/28 |
| TG1-5035 | Control | 2024/07/23 | 2024/08/06 | 2024/08/06 | 2024/09/05 | 2024/09/19 | 2024/10/04 |
| TG1-5038 | Control | 2024/07/26 | 2024/08/06 | 2024/08/06 | 2024/09/04 | 2024/09/25 | Breeding |
| TG1-5039 | Control | 2024/07/26 | 2024/08/06 | 2024/08/06 | . | . | 2024/08/28 |
| TG1-5043 | Control | 2024/07/30 | 2024/08/09 | 2024/08/09 | 2024/09/09 | 2024/09/25 | 2024/10/04 |
| TG1-5044 | Control | 2024/07/30 | 2024/08/09 | 2024/08/09 | . | . | 2024/08/15 |
| TG1-5046 | Control | 2024/07/30 | 2024/08/09 | 2024/08/09 | 2024/09/19 | 2024/10/02 | Breeding |
| TG1-5047 | Control | 2024/07/30 | 2024/08/09 | 2024/08/09 | 2024/09/13 | 2024/09/25 | Breeding |
| TG1-5048 | Control | 2024/07/30 | 2024/08/09 | 2024/08/09 | 2024/09/20 | 2024/10/02 | Breeding |
| TG1-5049 | Control | 2024/08/02 | 2024/08/16 | 2024/08/16 | 2024/09/27 | 2024/10/07 | Breeding |
| TG1-5050 | Control | 2024/08/02 | 2024/08/16 | 2024/08/16 | . | . | 2024/08/22 |
| TG1-5051 | Control | 2024/08/06 | 2024/08/16 | 2024/08/16 | 2024/09/17 | 2024/09/30 | 2024/10/04 |
| TG1-5052 | Control | 2024/08/02 | 2024/08/20 | 2024/08/20 | . | . | 2024/08/28 |
| TG1-5053 | Control | 2024/08/02 | 2024/08/20 | 2024/08/20 | . | . | 2024/08/28 |
| TG1-5054 | Control | 2024/08/02 | 2024/08/20 | 2024/08/20 | . | . | 2024/08/23 |
| TG1-5056 | Control | 2024/08/06 | 2024/08/20 | 2024/08/20 | . | . | 2024/08/28 |
| TG1-5057 | Control | 2024/08/06 | 2024/08/20 | 2024/08/20 | . | . | 2024/08/28 |
| TG1-5059 | Control | 2024/08/06 | 2024/08/20 | 2024/08/20 | . | . | 2024/08/23 |
| TG1-5060 | Control | 2024/08/09 | 2024/08/20 | 2024/08/20 | . | . | 2024/08/28 |
| TG1-5061 | Control | 2024/08/09 | 2024/08/20 | 2024/08/20 | 2024/09/20 | 2024/10/03 | Breeding |
| TG1-5062 | Control | 2024/08/09 | 2024/08/20 | 2024/08/20 | 2024/09/25 | 2024/10/07 | Breeding |
| TG1-5063 | Control | 2024/08/09 | 2024/08/20 | 2024/08/20 | 2024/09/30 | 2024/10/10 | Breeding |
| TG1-5064 | Control | 2024/08/09 | 2024/08/20 | 2024/08/20 | . | . | 2024/08/28 |
| TG1-5065 | Control | 2024/08/09 | 2024/08/20 | 2024/08/20 | . | . | 2024/08/28 |
| TG1-5066 | Control | 2024/08/09 | 2024/08/20 | 2024/08/20 | . | . | 2024/09/02 |
| TG1-5067 | Control | 2024/08/09 | 2024/08/20 | 2024/08/20 | 2024/09/25 | 2024/10/07 | Breeding |
| TG1-5068 | Control | 2024/08/09 | 2024/08/20 | 2024/08/20 | 2024/09/25 | 2024/10/02 | Breeding |
| TG1-5069 | Control | 2024/08/09 | 2024/08/20 | 2024/08/20 | . | . | 2024/09/13 |
| TG1-5070 | Control | 2024/08/09 | 2024/08/20 | 2024/08/20 | 2024/09/25 | 2024/10/10 | Breeding |
| TG1-5071 | Control | 2024/08/09 | 2024/08/20 | 2024/08/20 | 2024/09/25 | 2024/10/07 | Breeding |
| TG1-5072 | Control | 2024/08/09 | 2024/08/20 | 2024/08/20 | . | . | 2024/08/23 |
| TG1-5073 | Control | 2024/08/09 | 2024/08/20 | 2024/08/20 | . | . | 2024/08/28 |
| TG1-5074 | Control | 2024/08/09 | 2024/08/20 | 2024/08/20 | 2024/09/25 | 2024/10/02 | Breeding |
| TG1-5075 | Control | 2024/08/09 | 2024/08/20 | 2024/08/20 | . | . | 2024/08/28 |
| TG1-5076 | Control | 2024/08/09 | 2024/08/20 | 2024/08/20 | 2024/09/25 | . | 2024/10/07 |
| TG1-5078 | Control | 2024/08/09 | 2024/08/20 | 2024/08/20 | . | . | 2024/08/23 |
| TG1-5079 | Control | 2024/08/09 | 2024/08/20 | 2024/08/20 | . | . | 2024/08/22 |
| TG1-5080 | Control | 2024/08/09 | 2024/08/20 | 2024/08/20 | . | . | 2024/08/28 |
| TG1-5081 | Control | 2024/08/09 | 2024/08/20 | 2024/08/20 | . | . | 2024/08/23 |
| TG1-5082 | Control | 2024/08/09 | 2024/08/20 | 2024/08/20 | . | . | 2024/08/23 |
| TG1-5083 | Control | 2024/08/09 | 2024/08/20 | 2024/08/20 | . | . | 2024/08/28 |
| TG1-5084 | Control | 2024/08/16 | 2024/08/27 | 2024/08/27 | 2024/09/30 | 2024/10/17 | Breeding |
| TG1-5085 | Control | 2024/08/16 | 2024/08/27 | 2024/08/27 | . | . | 2024/09/02 |
| TG1-5086 | Control | 2024/08/16 | 2024/08/27 | 2024/08/27 | . | . | 2024/09/03 |
| TG1-5087 | Control | 2024/08/16 | 2024/08/27 | 2024/08/27 | . | . | 2024/09/27 |
| TG1-5088 | Control | 2024/08/16 | 2024/08/27 | 2024/08/27 | 2024/09/30 | 2024/10/17 | Breeding |
| TG1-5089 | Control | 2024/08/16 | 2024/08/27 | 2024/08/27 | . | . | 2024/09/13 |
| TG1-5090 | Control | 2024/08/16 | 2024/08/27 | 2024/08/27 | . | . | 2024/09/04 |
| TG1-5091 | Control | 2024/08/16 | 2024/08/27 | 2024/08/27 | . | . | 2024/09/02 |
| TG1-5092 | Control | 2024/08/16 | 2024/08/27 | 2024/08/27 | 2024/09/30 | 2024/10/11 | Breeding |
| TG1-5093 | Control | 2024/08/16 | 2024/08/27 | 2024/08/27 | . | . | 2024/09/13 |
| TG1-5094 | Control | 2024/08/16 | 2024/08/27 | 2024/08/27 | . | . | 2024/09/02 |
| TG1-5095 | Control | 2024/08/16 | 2024/08/27 | 2024/08/27 | . | . | 2024/09/03 |
| TG1-5096 | Control | 2024/08/16 | 2024/08/27 | 2024/08/27 | 2024/10/07 | 2024/10/21 | Breeding |
| TG1-5097 | Control | 2024/08/16 | 2024/08/27 | 2024/08/27 | . | . | 2024/09/02 |
| TG1-5098 | Control | 2024/08/16 | 2024/08/27 | 2024/08/27 | . | . | 2024/09/02 |
| TG1-5099 | Control | 2024/08/16 | 2024/08/27 | 2024/08/27 | . | . | 2024/09/02 |
| TG1-5101 | Control | 2024/08/16 | 2024/08/27 | 2024/08/27 | . | . | 2024/09/02 |
| TG1-5102 | Control | 2024/08/16 | 2024/08/27 | 2024/08/27 | . | . | 2024/09/02 |
| TG1-5103 | Control | 2024/08/16 | 2024/08/27 | 2024/08/27 | . | . | 2024/09/02 |
| TG1-5104 | Control | 2024/08/16 | 2024/08/27 | 2024/08/27 | 2024/09/30 | 2024/10/17 | Breeding |
| TG1-5105 | Control | 2024/08/16 | 2024/08/27 | 2024/08/27 | 2024/09/30 | 2024/10/11 | Breeding |
| TG1-5106 | Control | 2024/08/16 | 2024/08/27 | 2024/08/27 | 2024/10/04 | 2024/10/21 | Breeding |
| TG1-5107 | Control | 2024/08/16 | 2024/08/27 | 2024/08/27 | 2024/09/30 | 2024/10/10 | Breeding |
| TG1-5108 | Control | 2024/08/16 | 2024/08/27 | 2024/08/27 | . | . | 2024/09/02 |
| TG1-5109 | Control | 2024/08/16 | 2024/08/27 | 2024/08/27 | . | . | 2024/09/06 |
| TG1-5110 | Control | 2024/08/16 | 2024/08/27 | 2024/08/27 | . | . | 2024/09/02 |
| TG1-5111 | Control | 2024/08/16 | 2024/08/27 | 2024/08/27 | . | . | 2024/09/02 |
| TG1-5112 | Control | 2024/08/16 | 2024/08/27 | 2024/08/27 | . | . | 2024/09/02 |
| TG1-5113 | Control | 2024/08/16 | 2024/08/27 | 2024/08/27 | 2024/10/07 | 2024/10/28 | Breeding |
| TG1-5116 | Control | 2024/08/16 | 2024/08/27 | 2024/08/27 | . | . | 2024/09/02 |
| TG1-5117 | Control | 2024/08/16 | 2024/09/03 | 2024/09/03 | . | . | 2024/09/13 |
| TG1-5118 | Control | 2024/08/20 | 2024/09/03 | 2024/09/03 | 2024/10/07 | 2024/10/21 | Breeding |
| TG1-5119 | Control | 2024/08/20 | 2024/09/03 | 2024/09/03 | 2024/10/11 | 2024/10/24 | 2024/11/06 |
| TG1-5120 | Control | 2024/08/20 | 2024/09/03 | 2024/09/03 | . | . | 2024/09/06 |
| TG1-5121 | Control | 2024/08/20 | 2024/09/03 | 2024/09/03 | 2024/10/07 | 2024/10/21 | Breeding |
| TG1-5124 | Control | 2024/08/23 | 2024/09/03 | 2024/09/03 | . | . | 2024/09/13 |
| TG1-5125 | Control | 2024/08/23 | 2024/09/03 | 2024/09/03 | . | . | 2024/09/13 |
| TG1-5126 | Control | 2024/08/23 | 2024/09/03 | 2024/09/03 | 2024/10/03 | 2024/10/17 | Breeding |
| TG1-5129 | Control | 2024/08/23 | 2024/09/03 | 2024/09/03 | . | . | 2024/09/06 |
| TG1-5131 | Control | 2024/08/23 | 2024/09/03 | 2024/09/03 | 2024/10/16 | 2024/10/28 | Breeding |
| TG1-5132 | Control | 2024/08/23 | 2024/09/03 | 2024/09/03 | 2024/10/11 | 2024/10/28 | Breeding |
| TG1-5133 | Control | 2024/08/23 | 2024/09/03 | 2024/09/03 | 2024/10/07 | 2024/10/21 | Breeding |
| TG1-5134 | Control | 2024/08/23 | 2024/09/03 | 2024/09/03 | 2024/10/03 | . | 2025/10/10 |
| TG1-5135 | Control | 2024/08/23 | 2024/09/03 | 2024/09/03 | 2024/10/07 | 2024/10/21 | Breeding |
| TG1-5136 | Control | 2024/08/23 | 2024/09/03 | 2024/09/03 | 2024/10/03 | 2024/10/17 | Breeding |
| TG1-5139 | Control | 2024/08/23 | 2024/09/03 | 2024/09/03 | 2024/10/03 | 2024/10/17 | Breeding |
| TG1-5141 | Control | 2024/08/23 | 2024/09/03 | 2024/09/03 | 2024/10/04 | 2024/10/21 | Breeding |
| TG1-5142 | Control | 2024/08/23 | 2024/09/03 | 2024/09/03 | . | . | 2024/09/17 |
| TG1-5143 | Control | 2024/08/23 | 2024/09/03 | 2024/09/03 | . | . | 2024/09/17 |
| TG1-5144 | Control | 2024/08/23 | 2024/09/03 | 2024/09/03 | 2024/10/04 | 2024/10/13 | Breeding |
| TG1-5145 | Control | 2024/08/23 | 2024/09/03 | 2024/09/03 | . | . | 2024/09/06 |
| TG1-5146 | Control | 2024/08/23 | 2024/09/03 | 2024/09/03 | 2024/10/07 | 2024/10/21 | Breeding |
| TG1-5148 | Control | 2024/08/23 | 2024/09/03 | 2024/09/03 | 2024/10/04 | 2024/10/17 | Breeding |
| TG1-5149 | Control | 2024/08/23 | 2024/09/03 | 2024/09/03 | 2024/10/07 | 2024/10/28 | Breeding |
| TG1-5150 | Control | 2024/08/23 | 2024/09/03 | 2024/09/03 | . | . | 2024/09/06 |
| TG1-5151 | Control | 2024/08/27 | 2024/09/06 | 2024/09/06 | . | . | 2024/09/06 |
| TG1-5152 | Control | 2024/08/27 | 2024/09/06 | 2024/09/06 | . | . | 2024/09/06 |
| TG1-5154 | Control | 2024/08/27 | 2024/09/06 | 2024/09/06 | 2024/10/08 | 2024/10/28 | Breeding |
| TG1-5155 | Control | 2024/08/27 | 2024/09/06 | 2024/09/06 | 2024/10/11 | . | 2025/01/24 |
| TG1-5156 | Control | 2024/08/30 | 2024/09/10 | 2024/09/10 | . | . | 2024/09/13 |
| TG1-5157 | Control | 2024/08/30 | 2024/09/10 | 2024/09/10 | . | . | 2024/09/17 |
| TG1-5158 | Control | 2024/08/30 | 2024/09/10 | 2024/09/10 | . | . | 2024/09/17 |
| TG1-5159 | Control | 2024/08/30 | 2024/09/10 | 2024/09/10 | . | . | 2024/09/17 |
| TG1-5160 | Control | 2024/08/30 | 2024/09/10 | 2024/09/10 | . | . | 2024/09/17 |
| TG1-5161 | Control | 2024/08/30 | 2024/09/10 | 2024/09/10 | . | . | 2024/09/25 |
| TG1-5162 | Control | 2024/08/30 | 2024/09/10 | 2024/09/10 | . | . | 2024/09/19 |
| TG1-5163 | Control | 2024/08/30 | 2024/09/10 | 2024/09/10 | . | . | 2024/09/17 |
| TG1-5164 | Control | 2024/08/30 | 2024/09/10 | 2024/09/10 | . | . | 2024/09/17 |
| TG1-5165 | Control | 2024/08/30 | 2024/09/10 | 2024/09/10 | 2024/10/21 | . | no data |
| TG1-5166 | Control | 2024/08/30 | 2024/09/10 | 2024/09/10 | 2024/10/11 | 2024/10/28 | Breeding |
| TG1-5167 | Control | 2024/08/30 | 2024/09/10 | 2024/09/10 | . | . | 2024/09/17 |
| TG1-5168 | Control | 2024/08/30 | 2024/09/10 | 2024/09/10 | . | . | 2024/09/17 |
| TG1-5169 | Control | 2024/08/30 | 2024/09/10 | 2024/09/10 | . | . | 2024/09/17 |
| TG1-5172 | Control | 2024/08/30 | 2024/09/10 | 2024/09/10 | . | . | 2024/09/17 |
| TG1-5174 | Control | 2024/08/30 | 2024/09/10 | 2024/09/10 | 2024/10/16 | 2024/10/28 | Breeding |
| TG1-5175 | Control | 2024/08/30 | 2024/09/10 | 2024/09/10 | . | . | 2024/09/17 |
| TG1-5177 | Control | 2024/08/30 | 2024/09/10 | 2024/09/10 | 2024/10/18 | 2024/10/30 | Breeding |
| TG1-5178 | Control | 2024/08/30 | 2024/09/10 | 2024/09/10 | . | . | 2024/09/17 |
| TG1-5179 | Control | 2024/08/30 | 2024/09/10 | 2024/09/10 | 2024/10/11 | 2024/10/28 | Breeding |
| TG1-5180 | Control | 2024/08/30 | 2024/09/10 | 2024/09/10 | 2024/10/11 | 2024/10/24 | Breeding |
| TG1-5181 | Control | 2024/08/30 | 2024/09/10 | 2024/09/10 | 2024/10/16 | 2024/10/30 | Breeding |
| TG1-5182 | Control | 2024/08/30 | 2024/09/10 | 2024/09/10 | . | . | 2024/09/17 |
| TG1-5183 | Control | 2024/08/30 | 2024/09/10 | 2024/09/10 | 2024/10/16 | 2024/10/28 | Breeding |
| TG1-5184 | Control | 2024/08/30 | 2024/09/10 | 2024/09/10 | . | . | 2024/09/17 |
| TG1-5185 | Control | 2024/08/30 | 2024/09/10 | 2024/09/10 | . | . | 2024/09/17 |
| TG1-5186 | Control | 2024/08/30 | 2024/09/10 | 2024/09/10 | 2024/10/10 | 2024/10/24 | Breeding |
| TG1-5187 | Control | 2024/08/30 | 2024/09/10 | 2024/09/10 | . | . | 2024/09/17 |
| TG1-5188 | Control | 2024/08/30 | 2024/09/10 | 2024/09/10 | 2024/10/11 | 2024/10/24 | 2024/11/06 |
| TG1-5201 | Control | 2024/09/03 | 2024/09/13 | 2024/09/13 | . | . | 2024/10/08 |
| TG1-5202 | Control | 2024/09/03 | 2024/09/13 | 2024/09/13 | . | . | 2024/09/17 |
| TG1-5203 | Control | 2024/09/03 | 2024/09/13 | 2024/09/13 | 2024/10/16 | 2024/10/28 | Breeding |
| TG1-5204 | Control | 2024/09/03 | 2024/09/13 | 2024/09/13 | . | . | 2024/09/25 |
| TG1-5205 | Control | 2024/09/03 | 2024/09/13 | 2024/09/13 | 2024/10/16 | 2024/10/28 | Breeding |
| TG1-5206 | Control | 2024/09/03 | 2024/09/13 | 2024/09/13 | . | . | 2024/09/19 |
| TG1-5207 | Control | 2024/09/03 | 2024/09/13 | 2024/09/13 | 2024/10/21 | . | no data |
| TG1-5208 | Control | 2024/09/03 | 2024/09/13 | 2024/09/13 | 2024/10/21 | 2024/11/05 | Breeding |
| TG1-5210 | Control | 2024/09/03 | 2024/09/13 | 2024/09/13 | 2024/10/16 | 2024/10/30 | Breeding |
| TG1-5211 | Control | 2024/09/03 | 2024/09/13 | 2024/09/13 | 2024/10/21 | . | 2024/11/26 |
| TG1-5212 | Control | 2024/09/03 | 2024/09/13 | 2024/09/13 | . | . | 2024/10/22 |
| TG1-5213 | Control | 2024/09/03 | 2024/09/13 | 2024/09/13 | 2024/10/28 | 2024/11/11 | Breeding |
| TG1-5214 | Control | 2024/09/03 | 2024/09/13 | 2024/09/13 | . | . | 2024/09/19 |
| TG1-5215 | Control | 2024/09/03 | 2024/09/13 | 2024/09/13 | 2024/10/21 | 2024/11/05 | Breeding |
| TG1-5216 | Control | 2024/09/03 | 2024/09/13 | 2024/09/13 | . | . | 2024/09/19 |
| TG1-5217 | Control | 2024/09/03 | 2024/09/13 | 2024/09/13 | . | . | 2024/09/25 |
| TG1-5218 | Control | 2024/09/03 | 2024/09/13 | 2024/09/13 | . | . | 2024/09/19 |
| TG1-5219 | Control | 2024/09/03 | 2024/09/13 | 2024/09/13 | 2024/10/16 | 2024/10/30 | Breeding |
| TG1-5220 | Control | 2024/09/03 | 2024/09/13 | 2024/09/13 | . | . | 2024/09/25 |
| TG1-5221 | Control | 2024/09/03 | 2024/09/13 | 2024/09/13 | 2024/11/18 | 2024/11/27 | Breeding |
| TG1-5222 | Control | 2024/09/03 | 2024/09/13 | 2024/09/17 | 2024/10/21 | . | no data |
| TG1-5223 | Control | 2024/09/03 | 2024/09/13 | 2024/09/17 | 2024/10/28 | . | no data |
| TG1-5224 | Control | 2024/09/03 | 2024/09/13 | 2024/09/17 | 2024/10/18 | 2024/10/30 | Breeding |
| TG1-5225 | Control | 2024/09/06 | 2024/09/17 | 2024/09/17 | . | . | 2024/10/30 |
| TG1-5226 | Control | 2024/09/06 | 2024/09/17 | 2024/09/17 | 2024/10/21 | . | no data |
| TG1-5227 | Control | 2024/09/06 | 2024/09/17 | 2024/09/17 | . | . | 2024/10/30 |
| TG1-5228 | Control | 2024/09/06 | 2024/09/17 | 2024/09/17 | . | . | 2024/10/15 |
| TG1-5229 | Control | 2024/09/06 | 2024/09/17 | 2024/09/17 | 2024/10/30 | 2024/11/05 | Breeding |
| TG1-5230 | Control | 2024/09/06 | 2024/09/17 | 2024/09/17 | 2024/10/18 | 2024/10/30 | Breeding |
| TG1-5232 | Control | 2024/09/06 | 2024/09/17 | 2024/09/17 | . | . | 2024/10/07 |
| TG1-5233 | Control | 2024/09/06 | 2024/09/17 | 2024/09/17 | . | . | 2024/11/25 |
| TG1-5234 | Control | 2024/09/06 | 2024/09/17 | 2024/09/17 | 2024/10/18 | 2024/11/01 | Breeding |
| TG1-5236 | Control | 2024/09/06 | 2024/09/17 | 2024/09/17 | 2024/10/18 | 2024/10/28 | Breeding |
| TG1-5237 | Control | 2024/09/06 | 2024/09/17 | 2024/09/17 | 2024/10/18 | 2024/10/30 | Breeding |
| TG1-5238 | Control | 2024/09/06 | 2024/09/17 | 2024/09/17 | 2024/10/21 | 2024/11/05 | Breeding |
| TG1-5239 | Control | 2024/09/06 | 2024/09/17 | 2024/09/17 | 2024/10/21 | 2024/11/01 | Breeding |
| TG1-5240 | Control | 2024/09/06 | 2024/09/17 | 2024/09/17 | . | . | 2024/09/25 |
| TG1-5241 | Control | 2024/09/06 | 2024/09/17 | 2024/09/17 | . | . | 2024/09/25 |
| TG1-5242 | Control | 2024/09/06 | 2024/09/17 | 2024/09/17 | 2024/10/22 | . | no data |
| TG1-5243 | Control | 2024/09/06 | 2024/09/17 | 2024/09/17 | . | . | 2024/10/07 |
| TG1-5244 | Control | 2024/09/06 | 2024/09/17 | 2024/09/17 | 2024/10/21 | 2024/11/05 | no data |
| TG1-5245 | Control | 2024/09/06 | 2024/09/17 | 2024/09/17 | 2024/10/18 | 2024/10/30 | Breeding |
| TG1-5246 | Control | 2024/09/06 | 2024/09/17 | 2024/09/17 | 2024/10/18 | 2024/11/01 | Breeding |
| TG1-5247 | Control | 2024/09/06 | 2024/09/17 | 2024/09/17 | 2024/10/22 | . | no data |
| TG1-5248 | Control | 2024/09/06 | 2024/09/17 | 2024/09/17 | 2024/10/15 | 2024/10/28 | Breeding |
| TG1-5249 | Control | 2024/09/06 | 2024/09/17 | 2024/09/17 | 2024/10/22 | 2024/11/05 | Breeding |
| TG1-5250 | Control | 2024/09/06 | 2024/09/17 | 2024/09/17 | 2024/10/28 | 2024/11/11 | Breeding |
| TG1-5251 | Control | 2024/09/06 | 2024/09/17 | 2024/09/17 | 2024/10/28 | . | no data |
| TG1-5252 | Control | 2024/09/06 | 2024/09/17 | 2024/09/17 | . | . | 2024/10/01 |
| TG1-5253 | Control | 2024/09/06 | 2024/09/17 | 2024/09/17 | . | . | 2024/10/01 |
| TG1-5254 | Control | 2024/09/06 | 2024/09/17 | 2024/09/17 | 2024/10/28 | 2024/11/05 | no data |
| TG1-5255 | Control | 2024/09/06 | 2024/09/17 | 2024/09/17 | 2024/10/18 | 2024/11/01 | Breeding |
| TG1-5256 | Control | 2024/09/06 | 2024/09/17 | 2024/09/17 | 2024/10/16 | 2024/10/30 | Breeding |
| TG1-5257 | Control | 2024/09/06 | 2024/09/17 | 2024/09/17 | 2024/10/28 | 2024/11/11 | Breeding |
| TG1-5258 | Control | 2024/09/06 | 2024/09/17 | 2024/09/17 | . | . | 2024/10/30 |
| TG1-5259 | Control | 2024/09/06 | 2024/09/17 | 2024/09/17 | . | . | 2024/10/30 |
| TG1-5260 | Control | 2024/09/06 | 2024/09/17 | 2024/09/17 | 2024/10/18 | 2024/11/01 | Breeding |
| TG1-5261 | Control | 2024/09/06 | 2024/09/17 | 2024/09/17 | 2024/10/28 | 2024/11/11 | Breeding |
| TG1-5262 | Control | 2024/09/06 | 2024/09/17 | 2024/09/17 | . | . | 2024/10/07 |
| TG1-5263 | Control | 2024/09/06 | 2024/09/17 | 2024/09/17 | 2024/10/30 | 2024/11/14 | Breeding |
| TG1-5264 | Control | 2024/09/06 | 2024/09/17 | 2024/09/17 | . | . | 24/0/24 |
| TG1-5265 | Control | 2024/09/06 | 2024/09/17 | 2024/09/17 | . | . | 2024/10/01 |
| TG1-5268 | Control | 2024/09/06 | 2024/09/17 | 2024/09/17 | 2024/10/21 | . | no data |
| TG1-5269 | Control | 2024/09/06 | 2024/09/17 | 2024/09/17 | 2024/10/21 | no data | Breeding |
| TG1-5270 | Control | 2024/09/06 | 2024/09/17 | 2024/09/17 | . | . | 2024/10/15 |
| TG1-5271 | Control | 2024/09/06 | 2024/09/17 | 2024/09/17 | 2024/10/28 | . | no data |
| TG1-5274 | Control | 2024/09/10 | 2024/09/24 | 2024/09/24 | 2024/10/28 | 2024/11/11 | Breeding |
| TG1-5275 | Control | 2024/09/10 | 2024/09/24 | 2024/09/24 | 2024/10/28 | . | no data |
| TG1-5276 | Control | 2024/09/10 | 2024/09/24 | 2024/09/24 | 2024/10/30 | 2024/11/14 | Breeding |
| TG1-5278 | Control | 2024/09/10 | 2024/09/24 | 2024/09/24 | 2024/10/28 | 2024/11/11 | Breeding |
| TG1-5279 | Control | 2024/09/10 | 2024/09/24 | 2024/09/24 | 2024/10/30 | . | no data |
| TG1-5280 | Control | 2024/09/10 | 2024/09/24 | 2024/09/24 | . | . | 2024/10/08 |
| TG1-5281 | Control | 2024/09/10 | 2024/09/24 | 2024/09/24 | . | . | 2024/11/05 |
| TG1-5282 | Control | 2024/09/10 | 2024/09/24 | 2024/09/24 | 2024/10/28 | 2024/11/11 | Breeding |
| TG1-5283 | Control | 2024/09/10 | 2024/09/24 | 2024/09/24 | 2024/10/30 | 2024/11/11 | Breeding |
| TG1-5284 | Control | 2024/09/10 | 2024/09/24 | 2024/09/24 | . | . | 2024/12/02 |
| TG1-5285 | Control | 2024/09/10 | 2024/09/24 | 2024/09/24 | 2024/11/19 | 2024/11/25 | Breeding |
| TG1-5286 | Control | 2024/09/10 | 2024/09/24 | 2024/09/24 | 2024/10/30 | . | 2024/11/26 |
| TG1-5287 | Control | 2024/09/10 | 2024/09/24 | 2024/09/24 | . | . | 2024/11/19 |
| TG1-5288 | Control | 2024/09/10 | 2024/09/24 | 2024/09/24 | . | . | 2024/10/01 |
| TG1-5289 | Control | 2024/09/10 | 2024/09/24 | 2024/09/24 | 2024/10/28 | 2024/11/11 | Breeding |
| TG1-5290 | Control | 2024/09/10 | 2024/09/24 | 2024/09/24 | . | . | 2024/10/30 |
| TG1-5291 | Control | 2024/09/10 | 2024/09/24 | 2024/09/24 | 2024/11/01 | 2024/11/14 | Breeding |
| TG1-5292 | Control | 2024/09/10 | 2024/09/24 | 2024/09/24 | 2024/10/28 | no data | no data |
| TG1-5293 | Control | 2024/09/10 | 2024/09/24 | 2024/09/24 | . | . | 2024/10/07 |
| TG1-5294 | Control | 2024/09/10 | 2024/09/24 | 2024/09/24 | . | . | 2024/10/01 |
| TG1-5295 | Control | 2024/09/10 | 2024/09/24 | 2024/09/24 | 2024/11/05 | no data | Breeding |
| TG1-5296 | Control | 2024/09/10 | 2024/09/24 | 2024/09/24 | 2024/11/19 | 2024/11/27 | 2024/12/06 |
| TG1-5297 | Control | 2024/09/10 | 2024/09/24 | 2024/09/24 | 2024/10/30 | 2024/11/14 | Breeding |
| TG1-5298 | Control | 2024/09/10 | 2024/09/24 | 2024/09/24 | . | . | 2024/11/28 |
| TG1-5299 | Control | 2024/09/10 | 2024/09/24 | 2024/09/24 | 2024/10/28 | 2024/11/11 | Breeding |
| TG1-5300 | Control | 2024/09/10 | 2024/09/24 | 2024/09/24 | 2024/10/28 | 2024/11/11 | Breeding |
| TG1-5301 | Control | 2024/09/10 | 2024/09/24 | 2024/09/24 | . | . | 2024/11/06 |
| TG1-5302 | Control | 2024/09/10 | 2024/09/24 | 2024/09/24 | 2024/10/28 | 2024/11/11 | Breeding |
| TG1-5304 | Control | 2024/09/13 | 2024/09/24 | 2024/09/24 | . | . | 2024/10/07 |
| TG1-5305 | Control | 2024/09/13 | 2024/09/24 | 2024/09/24 | 2024/10/28 | 2024/11/11 | Breeding |
| TG1-5306 | Control | 2024/09/13 | 2024/09/24 | 2024/09/24 | 2024/10/28 | 2024/11/11 | Breeding |
| TG1-5307 | Control | 2024/09/13 | 2024/09/24 | 2024/09/24 | 2024/10/28 | 2024/11/11 | 2024/11/26 |
| TG1-5308 | Control | 2024/09/13 | 2024/09/24 | 2024/09/24 | 2024/11/01 | 2024/11/14 | Breeding |
| TG1-5309 | Control | 2024/09/13 | 2024/09/24 | 2024/09/24 | . | . | 2024/10/01 |
| TG1-5310 | Control | 2024/09/13 | 2024/09/24 | 2024/09/24 | . | . | 2024/10/01 |
| TG1-5311 | Control | 2024/09/13 | 2024/09/24 | 2024/09/24 | 2024/10/30 | 2024/11/11 | Breeding |
| TG1-5313 | Control | 2024/09/13 | 2024/09/24 | 2024/09/24 | 2024/10/30 | 2024/11/14 | Breeding |
| TG1-5314 | Control | 2024/09/13 | 2024/09/24 | 2024/09/24 | 2024/11/05 | no data | Breeding |
| TG1-5315 | Control | 2024/09/13 | 2024/09/24 | 2024/09/24 | 2024/11/01 | 2024/11/14 | 2024/11/26 |
| TG1-5316 | Control | 2024/09/13 | 2024/09/24 | 2024/09/24 | 2024/10/28 | 2024/11/11 | Breeding |
| TG1-5317 | Control | 2024/09/13 | 2024/09/24 | 2024/09/24 | 2024/10/30 | 2024/11/11 | Breeding |
| TG1-5318 | Control | 2024/09/13 | 2024/09/24 | 2024/09/24 | 2024/10/28 | 2024/11/11 | Breeding |
| TG1-5319 | Control | 2024/09/13 | 2024/09/24 | 2024/09/24 | 2024/10/28 | 2024/11/11 | Breeding |
| TG1-5320 | Control | 2024/09/13 | 2024/09/24 | 2024/09/24 | 2024/10/22 | 2024/11/05 | no data |
| TG1-5321 | Control | 2024/09/13 | 2024/09/24 | 2024/09/24 | 2024/10/30 | 2024/11/14 | 2024/11/26 |
| TG1-5322 | Control | 2024/09/13 | 2024/09/24 | 2024/09/24 | 2024/10/28 | . | 2024/11/26 |
| TG1-5323 | Control | 2024/09/13 | 2024/09/24 | 2024/09/24 | . | . | 2024/10/01 |
| TG1-5324 | Control | 2024/09/13 | 2024/09/24 | 2024/09/24 | 2024/11/05 | . | no data |
| TG1-5325 | Control | 2024/09/13 | 2024/09/24 | 2024/09/24 | 2024/10/30 | 2024/11/14 | Breeding |
| TG1-5326 | Control | 2024/09/13 | 2024/09/24 | 2024/09/24 | 2024/10/28 | 2024/11/11 | Breeding |
| TG1-5327 | Control | 2024/09/13 | 2024/09/24 | 2024/09/24 | 2024/10/30 | 2024/11/11 | Breeding |
| TG1-5328 | Control | 2024/09/13 | 2024/09/24 | 2024/09/24 | . | . | 2024/10/01 |
| TG1-5329 | Control | 2024/09/13 | 2024/09/24 | 2024/09/24 | 2024/10/30 | 2024/11/11 | Breeding |
| TG1-5330 | Control | 2024/09/13 | 2024/09/24 | 2024/09/24 | . | . | 2024/10/07 |
| TG1-5331 | Control | 2024/09/13 | 2024/09/24 | 2024/09/24 | 2024/11/01 | 2024/11/14 | Breeding |
| TG1-5332 | Control | 2024/09/17 | 2024/09/27 | 2024/09/27 | . | . | 2024/10/08 |
| TG1-5333 | Control | 2024/09/17 | 2024/09/27 | 2024/09/27 | . | . | 2024/10/07 |
| TG1-5334 | Control | 2024/09/17 | 2024/09/27 | 2024/09/27 | . | . | 2024/12/16 |
| TG1-5335 | Control | 2024/09/17 | 2024/09/27 | 2024/09/27 | 2024/10/31 | 2024/11/11 | Breeding |
| TG1-5336 | Control | 2024/09/17 | 2024/09/27 | 2024/09/27 | . | . | 2024/10/07 |
| TG1-5340 | Control | 2024/09/20 | 2024/10/01 | 2024/10/01 | no data | . | 2024/11/19 |
| TG1-5341 | Control | 2024/09/20 | 2024/10/01 | 2024/10/01 | 2024/11/19 | 2024/11/25 | 2024/12/06 |
| TG1-5342 | Control | 2024/09/20 | 2024/10/01 | 2024/10/01 | 2024/10/30 | . | 2024/11/14 |
| TG1-5343 | Control | 2024/09/20 | 2024/10/01 | 2024/10/01 | . | . | 2024/10/07 |
| TG1-5344 | Control | 2024/09/20 | 2024/10/01 | 2024/10/01 | no data | 2024/11/19 | Breeding |
| TG1-5345 | Control | 2024/09/20 | 2024/10/01 | 2024/10/01 | 2024/11/18 | 2024/11/27 | Breeding |
| TG1-5346 | Control | 2024/09/20 | 2024/10/01 | 2024/10/01 | . | . | 2024/10/07 |
| TG1-5347 | Control | 2024/09/20 | 2024/10/01 | 2024/10/01 | 2024/11/19 | 2024/11/27 | Breeding |
| TG1-5348 | Control | 2024/09/20 | 2024/10/01 | 2024/10/01 | 2024/11/19 | no data | no data |
| TG1-5349 | Control | 2024/09/20 | 2024/10/01 | 2024/10/01 | . | . | 2024/10/07 |
| TG1-5351 | Control | 2024/09/20 | 2024/10/01 | 2024/10/01 | 2024/11/19 | 2024/11/25 | Breeding |
| TG1-5352 | Control | 2024/09/20 | 2024/10/01 | 2024/10/01 | 2024/11/19 | no data | 2024/12/06 |
| TG1-5353 | Control | 2024/09/20 | 2024/10/01 | 2024/10/01 | . | . | 2024/10/07 |
| TG1-5354 | Control | 2024/09/20 | 2024/10/01 | 2024/10/01 | 2024/11/19 | 2024/11/27 | Breeding |
| TG1-5355 | Control | 2024/09/20 | 2024/10/01 | 2024/10/01 | 2024/11/18 | 2024/11/27 | Breeding |
| TG1-5356 | Control | 2024/09/20 | 2024/10/01 | 2024/10/01 | 2024/11/19 | . | 2024/11/25 |
| TG1-5357 | Control | 2024/09/20 | 2024/10/01 | 2024/10/01 | 2024/11/05 | . | 2024/11/26 |
| TG1-5358 | Control | 2024/09/20 | 2024/10/01 | 2024/10/01 | 2024/10/05 | no data | Breeding |
| TG1-5359 | Control | 2024/09/20 | 2024/10/01 | 2024/10/01 | . | . | 2024/10/07 |
| TG1-5360 | Control | 2024/09/20 | 2024/10/01 | 2024/10/01 | 2024/11/05 | no data | Breeding |
| TG1-5361 | Control | 2024/09/20 | 2024/10/01 | 2024/10/01 | . | . | 2024/10/07 |
| TG1-5362 | Control | 2024/09/20 | 2024/10/01 | 2024/10/01 | 2024/11/01 | 2024/11/14 | 2024/11/26 |
| TG1-5364 | Control | 2024/09/20 | 2024/10/01 | 2024/10/01 | 2024/10/31 | 2024/11/14 | Breeding |
| TG1-5366 | Control | 2024/09/20 | 2024/10/01 | 2024/10/01 | . | . | 2024/10/07 |
| TG1-5367 | Control | 2024/09/20 | 2024/10/01 | 2024/10/01 | 2024/11/12 | 2024/11/26 | 2024/12/06 |
| TG1-5368 | Control | 2024/09/20 | 2024/10/01 | 2024/10/01 | 2024/10/31 | 2024/11/14 | Breeding |
| TG1-5369 | Control | 2024/09/20 | 2024/10/01 | 2024/10/01 | . | . | 2024/10/08 |
| TG1-5370 | Control | 2024/09/20 | 2024/10/01 | 2024/10/01 | 2024/11/06 | no data | Breeding |
| TG1-5371 | Control | 2024/09/20 | 2024/10/01 | 2024/10/01 | . | . | 2024/10/07 |
| TG1-5372 | Control | 2024/09/20 | 2024/10/01 | 2024/10/01 | . | . | 2025/02/04 |
| TG1-5373 | Control | 2024/09/20 | 2024/10/01 | 2024/10/01 | . | . | 2024/10/22 |
| TG1-5380 | Control | 2024/09/24 | 2024/10/04 | 2024/10/04 | 2024/11/18 | 2024/11/27 | Breeding |
| TG1-5381 | Control | 2024/09/24 | 2024/10/08 | 2024/10/08 | . | . | 2024/11/05 |
| TG1-5382 | Control | 2024/09/24 | 2024/10/08 | 2024/10/08 | 2024/11/18 | 2024/11/27 | Breeding |
| TG1-5383 | Control | 2024/09/24 | 2024/10/08 | 2024/10/08 | 2024/11/19 | 2024/11/27 | 2024/12/06 |
| TG1-5384 | Control | 2024/09/24 | 2024/10/08 | 2024/10/08 | 2024/11/19 | 2024/11/27 | Breeding |
| TG1-5385 | Control | 2024/09/24 | 2024/10/08 | 2024/10/08 | . | . | 2024/10/15 |
| TG1-5386 | Control | 2024/09/24 | 2024/10/08 | 2024/10/08 | 2024/11/19 | 2024/12/03 | 2024/12/17 |
| TG1-5387 | Control | 2024/09/24 | 2024/10/08 | 2024/10/08 | 2024/11/19 | no data | 2024/12/06 |
| TG1-5388 | Control | 2024/09/24 | 2024/10/08 | 2024/10/08 | . | . | 2024/11/05 |
| TG1-5389 | Control | 2024/09/24 | 2024/10/08 | 2024/10/08 | 2024/11/18 | 2024/11/27 | Breeding |
| TG1-5390 | Control | 2024/09/24 | 2024/10/08 | 2024/10/08 | 2024/11/18 | 2024/11/27 | Breeding |
| TG1-5391 | Control | 2024/09/24 | 2024/10/08 | 2024/10/08 | 2024/11/18 | 2024/11/27 | Breeding |
| TG1-5392 | Control | 2024/09/24 | 2024/10/08 | 2024/10/08 | . | . | 2024/11/05 |
| TG1-5393 | Control | 2024/09/24 | 2024/10/08 | 2024/10/08 | 2024/11/18 | 2024/11/27 | Breeding |
| TG1-5394 | Control | 2024/09/24 | 2024/10/08 | 2024/10/08 | 2024/11/05 | no data | Breeding |
| TG1-5395 | Control | 2024/09/24 | 2024/10/08 | 2024/10/08 | 2024/11/05 | 2024/11/19 | 2024/11/29 |
| TG1-5396 | Control | 2024/09/24 | 2024/10/08 | 2024/10/08 | 2024/11/19 | 2024/11/27 | Breeding |
| TG1-5397 | Control | 2024/09/24 | 2024/10/08 | 2024/10/08 | . | . | 2024/10/15 |
| TG1-5398 | Control | 2024/09/24 | 2024/10/08 | 2024/10/08 | 2024/11/18 | 2024/11/27 | Breeding |
| TG1-5399 | Control | 2024/09/24 | 2024/10/08 | 2024/10/08 | . | . | 2024/12/16 |
| TG1-5401 | Control | 2024/09/24 | 2024/10/08 | 2024/10/08 | . | . | 2024/11/18 |
| TG1-5402 | Control | 2024/09/24 | 2024/10/08 | 2024/10/08 | . | . | 2024/11/06 |
| TG1-5404 | Control | 2024/09/24 | 2024/10/08 | 2024/10/08 | 2024/11/06 | 2024/11/19 | Breeding |
| TG1-5405 | Control | 2024/09/24 | 2024/10/08 | 2024/10/08 | no data | . | no data |
| TG1-5406 | Control | 2024/09/24 | 2024/10/08 | 2024/10/08 | 2024/11/18 | 2024/11/27 | Breeding |
| TG1-5407 | Control | 2024/09/24 | 2024/10/08 | 2024/10/08 | 2024/11/18 | 2024/11/27 | 2024/12/06 |
| TG1-5408 | Control | 2024/09/27 | 2024/10/08 | 2024/10/08 | 2024/11/19 | 2024/11/27 | Breeding |
| TG1-5409 | Control | 2024/09/27 | 2024/10/08 | 2024/10/08 | 2024/11/19 | 2024/12/03 | Breeding |
| TG1-5410 | Control | 2024/09/27 | 2024/10/08 | 2024/10/08 | 2024/11/19 | 2024/11/27 | Breeding |
| TG1-5411 | Control | 2024/09/27 | 2024/10/08 | 2024/10/08 | 2024/11/26 | no data | 2024/12/06 |
| TG1-5412 | Control | 2024/09/27 | 2024/10/08 | 2024/10/08 | . | . | 2024/10/15 |
| TG1-5414 | Control | 2024/09/27 | 2024/10/08 | 2024/10/08 | 2024/11/18 | 2024/11/27 | Breeding |
| TG1-5415 | Control | 2024/09/27 | 2024/10/08 | 2024/10/08 | 2024/11/19 | 2024/11/27 | Breeding |
| TG1-5416 | Control | 2024/09/27 | 2024/10/08 | 2024/10/08 | . | . | 2024/10/15 |
| TG1-5417 | Control | 2024/09/27 | 2024/10/08 | 2024/10/08 | 2024/11/18 | no data | 2024/12/06 |
| TG1-5418 | Control | 2024/09/27 | 2024/10/08 | 2024/10/08 | 2024/11/18 | 2024/11/27 | Breeding |
| TG1-5419 | Control | 2024/09/27 | 2024/10/08 | 2024/10/08 | 2024/11/18 | 2024/11/27 | Breeding |
| TG1-5420 | Control | 2024/09/27 | 2024/10/08 | 2024/10/08 | 2024/11/19 | 2024/12/03 | 2024/12/17 |
| TG1-5421 | Control | 2024/09/27 | 2024/10/08 | 2024/10/08 | 2024/11/18 | no data | no data |
| TG1-5422 | Control | 2024/09/27 | 2024/10/08 | 2024/10/08 | no data | 2024/11/19 | Breeding |
| TG1-5423 | Control | 2024/09/27 | 2024/10/08 | 2024/10/08 | . | . | 2024/12/16 |
| TG1-5424 | Control | 2024/09/27 | 2024/10/08 | 2024/10/08 | . | . | 2024/11/14 |
| TG1-5425 | Control | 2024/09/27 | 2024/10/08 | 2024/10/08 | 2024/11/18 | 2024/11/27 | Breeding |
| TG1-5426 | Control | 2024/09/27 | 2024/10/08 | 2024/10/08 | 2024/11/19 | . | no data |
| TG1-5428 | Control | 2024/09/27 | 2024/10/08 | 2024/10/08 | . | . | 2024/10/15 |
| TG1-5429 | Control | 2024/09/27 | 2024/10/08 | 2024/10/08 | . | . | 2024/11/26 |
| TG1-5430 | Control | 2024/09/27 | 2024/10/08 | 2024/10/08 | . | . | 2024/12/24 |
| TG1-5431 | Control | 2024/09/27 | 2024/10/08 | 2024/10/08 | 2024/11/18 | 2024/11/27 | Breeding |
| TG1-5432 | Control | 2024/09/27 | 2024/10/08 | 2024/10/08 | . | . | 2024/10/15 |
| TG1-5433 | Control | 2024/09/27 | 2024/10/08 | 2024/10/08 | 2024/11/20 | 2024/12/06 | Breeding |
| TG1-5437 | Control | 2024/10/01 | 2024/10/15 | 2024/10/15 | 2024/11/18 | 2024/11/27 | Breeding |
| TG1-5438 | Control | 2024/10/01 | 2024/10/15 | 2024/10/15 | 2024/11/19 | . | 2024/12/06 |
| TG1-5439 | Control | 2024/10/01 | 2024/10/15 | 2024/10/15 | 2024/11/26 | no data | 2024/12/06 |
| TG1-5440 | Control | 2024/10/01 | 2024/10/15 | 2024/10/15 | 2024/11/19 | 2024/12/03 | 2024/12/17 |
| TG1-5441 | Control | 2024/10/01 | 2024/10/15 | 2024/10/15 | . | . | 2024/10/22 |
| TG1-5442 | Control | 2024/10/01 | 2024/10/15 | 2024/10/15 | 2024/11/18 | 2024/12/03 | 2024/12/17 |
| TG1-5443 | Control | 2024/10/01 | 2024/10/15 | 2024/10/15 | . | . | 2024/10/22 |
| TG1-5444 | Control | 2024/10/01 | 2024/10/15 | 2024/10/15 | 2024/11/19 | 2024/12/03 | Breeding |
| TG1-5445 | Control | 2024/10/01 | 2024/10/15 | 2024/10/15 | 2024/11/19 | no data | no data |
| TG1-5446 | Control | 2024/10/01 | 2024/10/15 | 2024/10/15 | . | . | 2024/11/19 |
| TG1-5447 | Control | 2024/10/01 | 2024/10/15 | 2024/10/15 | . | . | 2024/10/22 |
| TG1-5448 | Control | 2024/10/01 | 2024/10/15 | 2024/10/15 | . | . | 2024/10/22 |
| TG1-5449 | Control | 2024/10/01 | 2024/10/15 | 2024/10/15 | . | . | 2024/01/17 |
| TG1-5450 | Control | 2024/10/01 | 2024/10/15 | 2024/10/15 | . | . | 2024/10/31 |
| TG1-5452 | Control | 2024/10/01 | 2024/10/15 | 2024/10/15 | . | . | 2024/11/19 |
| TG1-5454 | Control | 2024/10/01 | 2024/10/15 | 2024/10/15 | . | . | 2024/01/24 |
| TG1-5455 | Control | 2024/10/01 | 2024/10/15 | 2024/10/15 | . | . | 2024/11/25 |
| TG1-5456 | Control | 2024/10/01 | 2024/10/15 | 2024/10/15 | . | . | 2025/01/09 |
| TG1-5457 | Control | 2024/10/01 | 2024/10/15 | 2024/10/15 | . | . | 2024/10/24 |
| TG1-5458 | Control | 2024/10/04 | 2024/10/15 | 2024/10/15 | . | . | 2025/01/31 |
| TG1-5459 | Control | 2024/10/04 | 2024/10/15 | 2024/10/15 | 2024/11/25 | 2024/12/06 | Breeding |
| TG1-5460 | Control | 2024/10/04 | 2024/10/15 | 2024/10/15 | 2024/11/25 | 2024/12/06 | Breeding |
| TG1-5461 | Control | 2024/10/04 | 2024/10/15 | 2024/10/15 | 2024/11/25 | 2024/12/06 | 2025/01/31 |
| TG1-5462 | Control | 2024/10/04 | 2024/10/15 | 2024/10/15 | 2024/11/26 | 2024/12/06 | 2024/12/17 |
| TG1-5463 | Control | 2024/10/04 | 2024/10/15 | 2024/10/15 | 2024/11/19 | . | 2024/12/06 |
| TG1-5464 | Control | 2024/10/04 | 2024/10/15 | 2024/10/15 | 2024/11/25 | 2024/12/06 | 2024/12/17 |
| TG1-5465 | Control | 2024/10/04 | 2024/10/15 | 2024/10/15 | . | . | 2024/10/22 |
| TG1-5466 | Control | 2024/10/04 | 2024/10/15 | 2024/10/15 | 2024/11/19 | no data | 2024/12/06 |
| TG1-5467 | Control | 2024/10/04 | 2024/10/15 | 2024/10/15 | 2024/11/19 | 2024/12/06 | Breeding |
| TG1-5468 | Control | 2024/10/04 | 2024/10/15 | 2024/10/15 | 2024/11/20 | no data | no data |
| TG1-5469 | Control | 2024/10/04 | 2024/10/15 | 2024/10/15 | . | . | 2024/12/24 |
| TG1-5470 | Control | 2024/10/04 | 2024/10/15 | 2024/10/15 | . | . | 2024/12/25 |
| TG1-5471 | Control | 2024/10/04 | 2024/10/15 | 2024/10/15 | 2024/11/20 | 2024/12/03 | Breeding |
| TG1-5472 | Control | 2024/10/04 | 2024/10/15 | 2024/10/15 | 2024/11/20 | no data | no data |
| TG1-5474 | Control | 2024/10/04 | 2024/10/15 | 2024/10/15 | 2024/11/19 | 2024/12/03 | Breeding |
| TG1-5475 | Control | 2024/10/04 | 2024/10/15 | 2024/10/15 | . | . | 2024/10/24 |
| TG1-5476 | Control | 2024/10/04 | 2024/10/15 | 2024/10/15 | . | . | 2025/02/28 |
| TG1-5490 | Control | 2024/10/08 | 2024/10/22 | 2024/10/22 | 2024/11/26 | 2024/12/06 | 2024/12/17 |
| TG1-5491 | Control | 2024/10/08 | 2024/10/22 | 2024/10/22 | . | . | 2024/11/29 |
| TG1-5492 | Control | 2024/10/08 | 2024/10/22 | 2024/10/22 | . | . | 2025/01/20 |
| TG1-5493 | Control | 2024/10/08 | 2024/10/22 | 2024/10/22 | 2024/11/26 | 2024/12/06 | Breeding |
| TG1-5494 | Control | 2024/10/08 | 2024/10/22 | 2024/10/22 | . | . | 2024/10/28 |
| TG1-5495 | Control | 2024/10/08 | 2024/10/22 | 2024/10/22 | . | . | 2025/01/17 |
| TG1-5496 | Control | 2024/10/08 | 2024/10/22 | 2024/10/22 | . | . | 2025/01/24 |
| TG1-5498 | Control | 2024/10/08 | 2024/10/22 | 2024/10/22 | 2025/01/17 | no data | 2025/01/31 |
| TG1-5499 | Control | 2024/10/08 | 2024/10/22 | 2024/10/22 | 2024/11/20 | 2024/12/03 | Breeding |
| TG1-5500 | Control | 2024/10/08 | 2024/10/22 | 2024/10/22 | . | . | 2025/01/06 |
| TG1-5502 | Control | 2024/10/08 | 2024/10/22 | 2024/10/22 | . | . | 2025/01/20 |
| TG1-5503 | Control | 2024/10/08 | 2024/10/22 | 2024/10/22 | . | . | 2025/01/31 |
| TG1-5504 | Control | 2024/10/08 | 2024/10/22 | 2024/10/22 | no data | 2025/01/09 | 2025/01/24 |
| TG1-5505 | Control | 2024/10/08 | 2024/10/22 | 2024/10/22 | . | . | 2024/12/24 |
| TG1-5506 | Control | 2024/10/08 | 2024/10/22 | 2024/10/22 | 2024/12/12 | . | 2025/01/27 |
| TG1-5507 | Control | 2024/10/11 | 2024/10/22 | 2024/10/22 | 2024/12/02 | 2024/12/17 | Breeding |
| TG1-5509 | Control | 2024/10/11 | 2024/10/22 | 2024/10/22 | . | . | 2025/01/22 |
| TG1-5510 | Control | 2024/10/11 | 2024/10/22 | 2024/10/22 | . | . | Breeding |
| TG1-5511 | Control | 2024/10/11 | 2024/10/22 | 2024/10/22 | . | . | 2025/01/06 |
| TG1-5512 | Control | 2024/10/11 | 2024/10/22 | 2024/10/22 | 2025/01/09 | 2025/01/17 | no data |
| TG1-5514 | Control | 2024/10/11 | 2024/10/22 | 2024/10/22 | 2025/01/30 | . | no data |
| TG1-5515 | Control | 2024/10/18 | 2024/10/29 | 2024/10/29 | . | . | 2025/01/24 |
| TG1-5516 | Control | 2024/10/18 | 2024/10/29 | 2024/10/29 | 2024/12/02 | 2024/12/17 | 2024/12/30 |
| TG1-5517 | Control | 2024/10/18 | 2024/10/29 | 2024/10/29 | . | . | 2025/01/06 |
| TG1-5518 | Control | 2024/10/18 | 2024/10/29 | 2024/10/29 | . | . | 2024/11/25 |
| TG1-5519 | Control | 2024/10/18 | 2024/10/29 | 2024/10/29 | 2024/12/02 | 2024/12/17 | 2024/12/30 |
| TG1-5521 | Control | 2024/10/18 | 2024/10/29 | 2024/10/29 | 2024/12/05 | no data | no data |
| TG1-5522 | Control | 2024/10/18 | 2024/10/29 | 2024/10/29 | . | . | 2025/01/20 |
| TG1-5523 | Control | 2024/10/18 | 2024/10/29 | 2024/10/29 | . | . | 2025/01/09 |
| TG1-5524 | Control | 2024/10/18 | 2024/10/29 | 2024/10/29 | 2025/01/17 | 2025/01/31 | no data |
| TG1-5525 | Control | 2024/10/18 | 2024/10/29 | 2024/10/29 | . | . | 2024/11/25 |
| TG1-5534 | Control | 2024/10/22 | 2024/11/05 | 2024/11/05 | . | . | 2024/12/11 |
| TG1-5535 | Control | 2024/10/22 | 2024/11/05 | 2024/11/05 | . | . | 2025/01/22 |
| TG1-5536 | Control | 2024/10/22 | 2024/11/05 | 2024/11/05 | . | . | 2025/01/06 |
| TG1-5537 | Control | 2024/10/22 | 2024/11/05 | 2024/11/05 | no data | no data | 2024/12/06 |
| TG1-5538 | Control | 2024/10/22 | 2024/11/05 | 2024/11/05 | . | . | 2025/01/20 |
| TG1-5539 | Control | 2024/10/22 | 2024/11/05 | 2024/11/05 | . | . | 2024/11/18 |
| TG1-5540 | Control | 2024/10/22 | 2024/11/05 | 2024/11/05 | 2024/12/10 | no data | no data |
| TG1-5543 | Control | 2024/10/25 | 2024/11/05 | 2024/11/05 | 2024/12/12 | no data | no data |
| TG1-5544 | Control | 2024/10/25 | 2024/11/05 | 2024/11/05 | 2024/12/12 | no data | no data |
| TG1-5549 | Control | 2024/11/01 | 2024/11/15 | 2024/11/15 | . | . | 2024/12/05 |
| TG1-5550 | Control | 2024/11/12 | 2024/11/26 | 2024/11/26 | 2025/01/10 | no data | no data |
| TG1-5551 | Control | 2024/11/12 | 2024/11/26 | 2024/11/26 | . | . | 2024/11/29 |
| TG1-5552 | Control | 2024/11/12 | 2024/11/26 | 2024/11/26 | 2025/01/17 | no data | 2025/01/31 |
| TG1-5553 | Control | 2024/11/12 | 2024/11/26 | 2024/11/26 | . | . | 2024/12/05 |
| TG1-5555 | Control | 2024/11/12 | 2024/11/26 | 2024/11/26 | . | . | 2024/11/29 |
| TG1-5556 | Control | 2024/11/12 | 2024/11/26 | 2024/11/26 | no data | 2025/01/17 | Breeding |
| TG1-5558 | Control | 2024/11/15 | 2024/11/26 | 2024/11/26 | . | . | 2024/12/02 |
| TG1-5559 | Control | 2024/11/15 | 2024/11/26 | 2024/11/26 | 2025/01/17 | . | no data |
| TG1-5560 | Control | 2024/11/15 | 2024/11/26 | 2024/11/26 | . | . | 2025/01/21 |
| TG1-5561 | Control | 2024/11/15 | 2024/11/26 | 2024/11/26 | . | . | 2024/11/29 |
| TG1-5562 | Control | 2024/11/15 | 2024/11/26 | 2024/11/26 | . | . | 2025/01/06 |
| TG1-5564 | Control | 2024/11/15 | 2024/11/26 | 2024/11/26 | . | . | 2025/01/30 |
| TG1-5565 | Control | 2024/11/15 | 2024/11/26 | 2024/11/26 | no data | 2025/01/22 | 2025/01/31 |
| TG1-5566 | Control | 2024/11/08 | 2024/11/19 | 2024/11/19 | . | . | 2025/02/07 |
| TG1-5567 | Control | 2024/11/15 | 2024/11/26 | 2024/11/26 | 2025/01/21 | 2025/02/10 | 2025/02/10 |
| TG2-5001 | Control | 2024/07/19 | 2024/08/06 | 2024/08/06 | 2024/09/17 | 2024/09/27 | 2024/10/11 |
| TG2-5002 | Control | 2024/07/19 | 2024/08/06 | 2024/08/06 | 2024/09/10 | 2024/09/25 | Breeding |
| TG2-5019 | Control | 2024/08/23 | 2024/09/10 | 2024/09/10 | . | . | 2024/09/17 |
| TG2-5020 | Control | 2024/08/23 | 2024/09/10 | 2024/09/10 | . | . | 2024/09/26 |
| TG2-5021 | Control | 2024/08/23 | 2024/09/10 | 2024/09/10 | . | . | 2024/09/26 |
| TG2-5022 | Control | 2024/08/30 | 2024/09/10 | 2024/09/10 | . | . | 2024/10/17 |
| TG2-5023 | Control | 2024/08/30 | 2024/09/10 | 2024/09/10 | 2024/10/22 | 2024/11/11 | no data |
| TG2-5024 | Control | 2024/08/30 | 2024/09/10 | 2024/09/10 | . | . | 2024/09/26 |
| TG2-5025 | Control | 2024/08/30 | 2024/09/10 | 2024/09/10 | . | . | 2024/09/13 |
| TG2-5026 | Control | 2024/08/30 | 2024/09/10 | 2024/09/10 | . | . | 2024/09/19 |
| TG2-5027 | Control | 2024/08/30 | 2024/09/10 | 2024/09/10 | . | . | 2024/09/26 |
| TG2-5028 | Control | 2024/08/30 | 2024/09/10 | 2024/09/10 | . | . | 2024/11/14 |
| TG2-5029 | Control | 2024/08/30 | 2024/09/10 | 2024/09/10 | 2024/09/17 | 2024/09/30 | 2024/10/04 |
| TG2-5030 | Control | 2024/08/30 | 2024/09/10 | 2024/09/10 | . | . | 2024/09/26 |
| TG2-5031 | Control | 2024/08/30 | 2024/09/10 | 2024/09/10 | . | . | 2024/09/26 |
| TG2-5032 | Control | 2024/08/30 | 2024/09/10 | 2024/09/10 | 2024/10/17 | 2024/11/01 | 2024/11/11 |
| TG2-5033 | Control | 2024/08/30 | 2024/09/10 | 2024/09/10 | . | . | 2024/09/26 |
| TG2-5034 | Control | 2024/08/30 | 2024/09/10 | 2024/09/10 | . | . | 2024/09/13 |
| TG2-5035 | Control | 2024/08/30 | 2024/09/10 | 2024/09/10 | . | . | 2024/09/13 |
| TG1-T015 | Test | 2024/08/27 | 2024/09/06 | . | . | . | no data |
| TG1-T016 | Test | 2024/08/27 | 2024/09/06 | . | . | . | 2024/09/26 |
| TG1-T017 | Test | 2024/08/27 | 2024/09/06 | . | . | . | 2024/10/04 |
| TG1-T018 | Test | 2024/08/27 | 2024/09/06 | . | . | . | no data |
| TG1-T019 | Test | 2024/08/27 | 2024/09/06 | . | . | . | 2024/10/25 |
| TG1-T020 | Test | 2024/08/27 | 2024/09/06 | . | . | . | 2024/09/26 |
| TG1-T021 | Test | 2024/08/27 | 2024/09/06 | . | . | . | 2024/10/17 |
| TG1-J001 | Combination | 2024/11/26 | 2024/12/06 | 2025/01/08 | 2025/01/20 | . | 2025/02/10 |
| TG1-J002 | Combination | 2024/11/26 | 2024/12/06 | . | . | . | 2025/01/08 |
| TG1-J003 | Combination | 2024/11/26 | 2024/12/06 | 2025/01/08 | 2025/01/22 | . | 2025/02/26 |
| TG1-J004 | Combination | 2024/11/26 | 2024/12/06 | . | . | . | 2025/01/08 |
| TG1-J005 | Combination | 2024/11/26 | 2024/12/06 | . | . | . | 2025/01/08 |
| TG1-J006 | Combination | 2024/11/26 | 2024/12/06 | . | . | . | 2025/01/08 |
| TG1-J008 | Combination | 2024/11/26 | 2024/12/06 | . | . | . | 2025/01/08 |
| TG1-J009 | Combination | 2024/11/26 | 2024/12/06 | 2025/01/08 | . | . | 2025/01/24 |
| TG1-J010 | Combination | 2024/11/26 | 2024/12/06 | 2025/01/08 | 2025/01/20 | . | 2025/02/26 |
| TG1-J011 | Combination | 2024/11/26 | 2024/12/06 | . | . | . | 2025/01/08 |
| TG1-J012 | Combination | 2024/11/26 | 2024/12/06 | 2025/01/08 | 2025/01/17 | . | 2025/02/10 |
| TG1-J013 | Combination | 2024/11/26 | 2024/12/06 | 2025/01/08 | 2025/01/17 | . | 2025/02/26 |
| TG1-J014 | Combination | 2024/11/26 | 2024/12/06 | . | . | . | 2025/01/08 |
| TG1-J015 | Combination | 2024/11/26 | 2024/12/06 | . | . | . | 2025/01/08 |
| TG1-J016 | Combination | 2024/11/26 | 2024/12/06 | 2025/01/08 | 2025/01/17 | . | 2025/02/26 |
| TG1-J017 | Combination | 2024/11/26 | 2024/12/06 | 2025/01/08 | 2025/01/20 | . | 2025/02/26 |
| TG1-J018 | Combination | 2024/11/26 | 2024/12/06 | 2025/01/08 | 2025/01/17 | . | 2025/02/10 |
| TG1-J019 | Combination | 2024/11/26 | 2024/12/06 | 2025/01/08 | . | . | 2025/01/17 |
| TG1-J020 | Combination | 2024/11/26 | 2024/12/10 | 2025/01/08 | . | . | 2025/02/12 |
| TG1-J021 | Combination | 2024/11/26 | 2024/12/10 | 2025/01/08 | 2025/01/20 | . | 2025/02/26 |
| TG1-J022 | Combination | 2024/11/26 | 2024/12/10 | . | . | . | 2025/01/08 |
| TG1-J023 | Combination | 2024/11/26 | 2024/12/10 | . | . | . | 2025/01/08 |
| TG1-J025 | Combination | 2024/11/26 | 2024/12/10 | 2025/01/08 | 2025/01/20 | . | no data |
| TG1-J026 | Combination | 2024/11/26 | 2024/12/10 | . | . | . | 2025/01/08 |
| TG1-J027 | Combination | 2024/11/26 | 2024/12/10 | . | . | . | 2025/01/08 |
| TG1-J028 | Combination | 2024/11/26 | 2024/12/10 | . | . | . | 2025/01/08 |
| TG1-J029 | Combination | 2024/11/26 | 2024/12/10 | . | . | . | 2025/01/08 |
| TG1-J030 | Combination | 2024/11/26 | 2024/12/10 | . | . | . | 2025/01/08 |
| TG1-J031 | Combination | 2024/11/26 | 2024/12/10 | 2025/01/08 | 2025/01/17 | 2025/01/30 | 2025/02/10 |
| TG1-J032 | Combination | 2024/11/26 | 2024/12/10 | . | . | . | 2025/01/08 |
| TG1-J033 | Combination | 2024/11/26 | 2024/12/10 | 2025/01/08 | . | . | 2025/01/17 |
| TG1-J034 | Combination | 2024/11/26 | 2024/12/10 | 2025/01/08 | . | . | 2025/01/29 |
| TG1-J035 | Combination | 2024/11/26 | 2024/12/10 | . | . | . | 2025/01/08 |
| TG1-J036 | Combination | 2024/11/29 | 2024/12/13 | . | . | . | 2025/01/10 |
| TG1-J037 | Combination | 2024/11/29 | 2024/12/13 | 2025/01/10 | 2025/01/28 | . | no data |
| TG1-J038 | Combination | 2024/11/29 | 2024/12/13 | . | . | . | 2025/01/10 |
| TG1-J039 | Combination | 2024/11/29 | 2024/12/13 | . | . | . | 2025/01/10 |
| TG1-J040 | Combination | 2024/11/29 | 2024/12/13 | 2025/01/10 | 2025/01/24 | no data | 2025/02/10 |
| TG1-J041 | Combination | 2024/11/29 | 2024/12/13 | 2025/01/10 | 2025/01/20 | . | 2025/01/31 |
| TG1-J042 | Combination | 2024/11/29 | 2024/12/13 | . | . | . | 2025/01/10 |
| TG1-J043 | Combination | 2024/11/29 | 2024/12/13 | . | . | . | 2025/01/10 |
| TG1-J044 | Combination | 2024/11/29 | 2024/12/13 | . | . | . | 2025/01/10 |
| TG1-J045 | Combination | 2024/11/29 | 2024/12/13 | . | . | . | 2025/01/10 |
| TG1-J046 | Combination | 2024/11/29 | 2024/12/13 | 2025/01/10 | 2025/01/30 | . | no data |
| TG1-J047 | Combination | 2024/11/29 | 2024/12/13 | 2025/01/10 | 2025/01/24 | no data | 2025/02/10 |
| TG1-J048 | Combination | 2024/11/29 | 2024/12/13 | . | . | . | 2025/01/10 |
| TG1-J049 | Combination | 2024/11/29 | 2024/12/13 | . | . | . | 2025/01/10 |
| TG1-J050 | Combination | 2024/11/29 | 2024/12/13 | 2025/01/10 | 2025/01/20 | . | 2025/02/10 |
| TG1-J051 | Combination | 2024/11/29 | 2024/12/13 | . | . | . | 2025/01/10 |
| TG1-J052 | Combination | 2024/11/29 | 2024/12/13 | . | . | . | 2025/01/10 |
| TG1-J053 | Combination | 2024/11/29 | 2024/12/13 | . | . | . | 2025/01/10 |
| TG1-J054 | Combination | 2024/11/29 | 2024/12/13 | . | . | . | 2025/01/10 |
| TG1-J055 | Combination | 2024/11/29 | 2024/12/13 | 2025/01/10 | . | . | 2025/01/29 |
| TG1-J056 | Combination | 2024/11/29 | 2024/12/13 | 2025/01/10 | 2025/01/24 | . | 2025/02/10 |
| TG1-J057 | Combination | 2024/11/29 | 2024/12/13 | 2025/01/10 | 2025/01/24 | 2025/02/07 | no data |
| TG1-J058 | Combination | 2024/11/29 | 2024/12/13 | . | . | . | 2025/01/10 |
| TG1-J059 | Combination | 2024/11/29 | 2024/12/13 | 2025/01/10 | 2025/01/17 | 2025/01/30 | 2025/02/10 |
| TG1-J060 | Combination | 2024/11/29 | 2024/12/13 | 2025/01/10 | 2025/01/20 | no data | 2025/02/10 |
| TG1-J061 | Combination | 2024/11/29 | 2024/12/13 | . | . | . | 2025/01/10 |
| TG1-J062 | Combination | 2024/11/29 | 2024/12/13 | . | . | . | 2025/01/10 |
| TG1-J063 | Combination | 2024/11/29 | 2024/12/13 | . | . | . | 2025/01/10 |
| TG1-J064 | Combination | 2024/11/29 | 2024/12/13 | 2025/01/10 | 2025/01/17 | . | 2025/01/31 |
| TG1-J065 | Combination | 2024/11/29 | 2024/12/13 | . | . | . | 2025/01/10 |
| TG1-J066 | Combination | 2024/11/29 | 2024/12/13 | . | . | . | 2025/01/10 |
| TG1-J067 | Combination | 2024/11/29 | 2024/12/13 | 2025/01/10 | . | . | 2025/02/21 |
| TG1-J069 | Combination | 2024/12/03 | 2024/12/18 | . | . | . | 2025/01/10 |
| TG1-J070 | Combination | 2024/12/03 | 2024/12/18 | 2025/01/10 | 2025/01/24 | . | 2025/02/26 |
| TG1-J071 | Combination | 2024/12/03 | 2024/12/18 | . | . | . | 2025/01/10 |
| TG1-J072 | Combination | 2024/12/03 | 2024/12/18 | . | . | . | 2025/01/10 |
| TG1-J073 | Combination | 2024/12/03 | 2024/12/18 | . | . | . | 2025/01/10 |
| TG1-J074 | Combination | 2024/12/03 | 2024/12/18 | 2025/01/10 | . | . | 2025/01/17 |
| TG1-J075 | Combination | 2024/12/03 | 2024/12/18 | 2025/01/10 | . | . | 2025/01/17 |
| TG1-J076 | Combination | 2024/12/03 | 2024/12/18 | . | . | . | 2025/01/10 |
| TG1-J077 | Combination | 2024/12/03 | 2024/12/18 | . | . | . | 2025/01/10 |
| TG1-J078 | Combination | 2024/12/06 | 2024/12/20 | . | . | . | 2025/01/14 |
| TG1-J079 | Combination | 2024/12/06 | 2024/12/20 | . | . | . | 2025/01/14 |
| TG1-J080 | Combination | 2024/12/06 | 2024/12/20 | 2025/01/14 | . | . | 2025/01/20 |
| TG1-J081 | Combination | 2024/12/06 | 2024/12/20 | 2025/01/14 | 2025/01/28 | . | 2025/02/26 |
| TG1-J082 | Combination | 2024/12/06 | 2024/12/20 | . | . | . | 2025/01/14 |
| TG1-J083 | Combination | 2024/12/06 | 2024/12/20 | . | . | . | 2025/01/14 |
| TG1-J084 | Combination | 2024/12/06 | 2024/12/20 | . | . | . | 2025/01/14 |
| TG1-J085 | Combination | 2024/12/06 | 2024/12/20 | 2025/01/14 | 2025/01/30 | . | 2025/02/26 |
| TG1-J086 | Combination | 2024/12/06 | 2024/12/20 | . | . | . | 2025/01/14 |
| TG1-J087 | Combination | 2024/12/06 | 2024/12/20 | . | . | . | 2025/01/14 |
| TG1-J088 | Combination | 2024/12/06 | 2024/12/20 | . | . | . | 2025/01/14 |
| TG1-J089 | Combination | 2024/12/06 | 2024/12/20 | 2025/01/14 | . | . | 2025/01/28 |
| TG1-J090 | Combination | 2024/12/06 | 2024/12/20 | . | . | . | 2025/01/14 |
| TG1-J091 | Combination | 2024/12/06 | 2024/12/20 | 2025/01/14 | 2025/01/30 | 2025/02/10 | 2025/02/26 |
| TG1-J092 | Combination | 2024/12/06 | 2024/12/20 | 2025/01/14 | 2025/01/29 | . | 2025/02/26 |
| TG1-J093 | Combination | 2024/12/06 | 2024/12/20 | 2025/01/14 | . | . | 2025/01/24 |
| TG1-J094 | Combination | 2024/12/06 | 2024/12/20 | . | . | . | 2025/01/14 |
| TG1-J095 | Combination | 2024/12/06 | 2024/12/20 | . | . | . | 2025/01/14 |
| TG1-J096 | Combination | 2024/12/06 | 2024/12/20 | . | . | . | 2025/01/14 |
| TG1-J097 | Combination | 2024/12/06 | 2024/12/20 | 2025/01/14 | . | . | 2025/01/24 |
| TG1-J098 | Combination | 2024/12/06 | 2024/12/20 | 2025/01/14 | 2025/01/24 | no data | 2025/02/10 |
| TG1-J099 | Combination | 2024/12/06 | 2024/12/20 | 2025/01/14 | 2025/01/20 | 2025/02/10 | 2025/02/10 |
| TG1-J100 | Combination | 2024/12/06 | 2024/12/20 | 2025/01/14 | 2025/01/29 | . | 2025/02/26 |
| TG1-J108 | Combination | 2024/12/13 | 2024/12/27 | . | . | . | 2025/01/14 |
| TG1-J110 | Combination | 2024/12/13 | 2024/12/27 | . | . | . | 2025/01/14 |
| TG1-J112 | Combination | 2024/12/13 | 2024/12/27 | . | . | . | 2025/01/14 |
| TG1-J118 | Combination | 2024/12/13 | 2024/12/27 | . | . | . | 2025/01/14 |
| TG1-J120 | Combination | 2024/12/13 | 2024/12/27 | . | . | . | 2025/01/14 |
| TG1-J127 | Combination | 2024/12/27 | 2025/01/10 | . | . | . | 2025/01/23 |
| TG1-J129 | Combination | 2024/12/27 | 2025/01/10 | . | . | . | 2025/01/23 |
| TG1-J130 | Combination | 2024/12/27 | 2025/01/10 | . | . | . | 2025/01/23 |
| TG1-J131 | Combination | 2024/12/27 | 2025/01/10 | . | . | . | 2025/01/23 |
| TG1-J132 | Combination | 2024/12/27 | 2025/01/10 | . | . | . | 2025/01/23 |
| TG1-J134 | Combination | 2024/12/27 | 2025/01/10 | . | . | . | 2025/01/23 |
| TG1-J135 | Combination | 2024/12/27 | 2025/01/10 | . | . | . | 2025/01/23 |
| TG1-J136 | Combination | 2024/12/27 | 2025/01/10 | . | . | . | 2025/01/23 |
| TG2-J001 | Combination | 2024/11/29 | 2024/12/18 | . | . | . | 2025/01/15 |
| TG2-J002 | Combination | 2024/12/06 | 2024/12/20 | . | . | . | 2025/01/15 |
| TG2-J003 | Combination | 2024/12/06 | 2024/12/20 | 2025/01/15 | 2025/01/20 | . | 2025/01/31 |
| TG2-J004 | Combination | 2024/12/06 | 2024/12/20 | . | . | . | 2025/01/15 |
| TG2-J005 | Combination | 2024/12/13 | 2024/12/27 | . | . | . | 2025/01/15 |
| TG2-J007 | Combination | 2024/12/13 | 2024/12/27 | . | . | . | 2025/01/15 |
| TG2-J009 | Combination | 2024/12/13 | 2024/12/27 | . | . | . | 2025/01/15 |
| TG2-J010 | Combination | 2024/12/13 | 2024/12/27 | . | . | . | 2025/01/15 |
| TG2-J011 | Combination | 2024/12/13 | 2024/12/27 | . | . | . | 2025/01/15 |
| TG2-J012 | Combination | 2024/12/13 | 2024/12/27 | . | . | . | 2025/01/15 |
| TG2-J013 | Combination | 2024/12/13 | 2024/12/27 | . | . | . | 2025/01/15 |
| TG2-J015 | Combination | 2024/12/13 | 2024/12/27 | . | . | . | 2025/01/15 |
| TG2-J016 | Combination | 2024/12/13 | 2024/12/27 | . | . | . | 2025/01/15 |
| TG2-J017 | Combination | 2024/12/13 | 2024/12/27 | . | . | . | 2025/01/15 |
| TG2-J018 | Combination | 2024/12/13 | 2024/12/27 | . | . | . | 2025/01/15 |
| TG2-J019 | Combination | 2024/12/24 | 2025/01/07 | . | . | . | 2025/01/15 |
| TG2-J020 | Combination | 2024/12/24 | 2025/01/07 | . | . | . | 2025/01/15 |
| TG2-J021 | Combination | 2024/12/24 | 2025/01/07 | . | . | . | 2025/01/15 |
| TG2-J022 | Combination | 2024/12/24 | 2025/01/07 | . | . | . | 2025/01/15 |
| TG2-J025 | Combination | 2024/12/20 | 2025/01/07 | . | . | . | 2025/01/15 |
| TG2-J026 | Combination | 2024/12/20 | 2025/01/07 | . | . | . | 2025/01/15 |
| TG2-J027 | Combination | 2024/12/27 | 2025/01/10 | . | . | . | 2025/01/23 |
| TG2-J029 | Combination | 2024/12/27 | 2025/01/10 | . | . | . | 2025/01/23 |
| TG2-J030 | Combination | 2024/12/27 | 2025/01/10 | . | . | . | 2025/01/23 |
| TG2-J031 | Combination | 2024/12/27 | 2025/01/10 | . | . | . | 2025/01/23 |
| TG2-J032 | Combination | 2024/12/27 | 2025/01/10 | . | . | . | 2025/01/23 |
| TG2-J033 | Combination | 2024/12/27 | 2025/01/10 | . | . | . | 2025/01/23 |
| TG2-J034 | Combination | 2024/12/27 | 2025/01/10 | . | . | . | 2025/01/23 |
| TG2-J051 | Combination | 2024/12/29 | 2025/01/10 | . | . | . | 2025/01/28 |
| TG2-J053 | Combination | 2024/12/29 | 2025/01/10 | . | . | . | 2025/01/28 |
| TG2-J055 | Combination | 2024/12/29 | 2025/01/10 | . | . | . | 2025/01/28 |
| TG2-J056 | Combination | 2024/12/29 | 2025/01/10 | . | . | . | 2025/01/28 |
| TG2-J057 | Combination | 2024/12/29 | 2025/01/10 | . | . | . | 2025/01/28 |
| BxW-01 | Tube | 2023/10/10 | 2023/11/07 | . | . | . | 2024/02/20 |
| BxW-02 | Tube | 2023/10/10 | 2023/11/07 | . | . | . | 2023/12/19 |
| BxW-03 | Tube | 2023/10/10 | 2023/11/07 | . | . | . | 2023/12/26 |
| BxW-04 | Tube | 2023/10/13 | 2023/11/07 | . | . | . | 2023/11/21 |
| BxW-05 | Tube | 2023/10/13 | 2023/11/07 | . | . | . | No data |
| BxW-06 | Tube | 2023/10/13 | 2023/11/10 | . | . | . | 2023/12/26 |
| BxW-07 | Tube | 2023/10/13 | 2023/11/14 | . | 2024/1/9 | . | No data |
| BxW-08 | Tube | 2023/10/13 | 2023/11/17 | . | . | . | No data |
| BxW-09 | Tube | 2023/10/13 | 2023/11/17 | . | . | . | 2024/02/20 |
| BxW-10 | Tube | 2023/10/13 | 2023/11/21 | . | . | . | 2024/01/04 |
| BxW-11 | Tube | 2023/11/21 | 2023/12/22 | . | . | . | 2024/03/26 |
| BxW-12 | Tube | 2023/11/21 | 2023/12/22 | 2024/04/22 | 2024/05/07 | 2024/05/20 | Breedig |
| BxW-13 | Tube | 2023/12/12 | 2024/01/12 | 2024/04/22 | 2024/05/10 | . | No data |
| BxW-14 | Tube | 2023/12/14 | 2024/01/12 | . | . | . | 2024/04/05 |
| BxW-15 | Tube | 2023/12/12 | 2024/01/16 | . | . | . | 2024/04/05 |
| BxW-16 | Tube | 2023/12/12 | 2024/01/16 | . | . | . | 2024/01/30 |
| BxW-17 | Tube | 2023/12/12 | 2024/01/16 | 2024/04/22 | 2024/05/07 | 2024/05/21 | Breedig |
| BxW-18 | Tube | 2023/12/12 | 2024/01/26 | 2024/04/22 | 2024/05/07 | 2024/05/21 | Breedig |
| BxW-19 | Tube | 2023/12/12 | 2024/01/26 | . | . | . | 2024/02/20 |
| BxW-20 | Tube | 2023/12/26 | 2024/01/26 | . | . | . | 2024/03/19 |
| BxW-21 | Tube | 2023/12/12 | 2024/01/30 | . | . | . | 2024/03/26 |
| BxW-22 | Tube | 2023/12/14 | 2024/01/30 | . | . | . | 2024/03/26 |
| BxW-23 | Tube | 2023/12/28 | 2024/02/02 | . | . | . | 2024/03/26 |
| BxW-24 | Tube | 2023/12/26 | 2024/02/02 | 2024/04/22 | 2024/05/07 | 2024/05/21 | Breedig |
| BxW-25 | Tube | 2023/12/26 | 2024/02/02 | 2024/04/22 | 2024/05/07 | 2024/05/20 | Breedig |
| BxW-26 | Tube | 2023/12/12 | 2024/01/23 | 2024/04/22 | 2024/05/07 | 2024/05/21 | Breedig |
| BxW-27 | Tube | 2023/12/12 | 2024/01/23 | 2024/04/22 | 2024/05/07 | 2024/05/22 | Breedig |
| WxB-01 | Tube | 2023/10/02 | 2023/10/27 | . | . | . | 2024/02/06 |
| WxB-02 | Tube | 2023/10/02 | 2023/10/27 | . | . | . | 2023/11/10 |
| WxB-03 | Tube | 2023/10/06 | 2023/10/27 | . | 2023/12/01 | . | 2024/01/05 |
| WxB-04 | Tube | 2023/10/06 | 2023/10/27 | 2024/04/22 | 2024/05/16 | 2024/5/28 | Breeding |
| WxB-05 | Tube | 2023/10/06 | 2023/10/27 | . | . | . | 2023/12/12 |
| WxB-06 | Tube | 2023/10/06 | 2023/10/27 | . | . | . | 2024/1/4 |
| WxB-07 | Tube | 2023/10/06 | 2023/10/27 | 2024/4/22 | . | . | 2024/5/13 |
| WxB-08 | Tube | 2023/10/02 | 2023/11/02 | . | . | . | 2023/11/13 |
| WxB-09 | Tube | 2023/10/02 | 2023/11/02 | 2024/4/22 | 2024/05/07 | 2024/05/20 | Breeding |
| WxB-10 | Tube | 2023/10/02 | 2023/11/02 | . | . | . | No data |
| WxB-11 | Tube | 2023/10/06 | 2023/11/07 | . | . | . | 2023/12/19 |
| WxB-12 | Tube | 2023/10/10 | 2023/11/07 | . | 2023/12/12 | . | No data |
| WxB-13 | Tube | 2023/10/13 | 2023/11/07 | 2024/04/22 | 2024/05/20 | . | No data |
| WxB-14 | Tube | 2023/10/13 | 2023/11/07 | . | . | . | 2023/11/21 |
| WxB-15 | Tube | 2023/10/02 | 2023/11/10 | . | . | . | 2023/12/19 |
| WxB-16 | Tube | 2023/10/06 | 2023/11/10 | . | 2023/12/12 | 2023/12/26 | Breeding |
| WxB-17 | Tube | 2023/10/06 | 2023/11/10 | 2024/04/22 | . | . | No data |
| WxB-18 | Tube | 2023/10/10 | 2023/11/10 | 2024/04/22 | 2024/05/07 | . | 2024/05/27 |
| WxB-19 | Tube | 2023/10/10 | 2023/11/10 | . | . | . | 2023/12/19 |
| WxB-20 | Tube | 2023/10/13 | 2023/11/10 | . | . | . | 2023/11/24 |
| WxB-21 | Tube | 2023/10/16 | 2023/11/10 | . | . | . | 2023/12/15 |
| WxB-22 | Tube | 2023/10/16 | 2023/11/10 | . | . | . | 2023/11/24 |
| WxB-23 | Tube | 2023/10/10 | 2023/11/14 | . | . | . | 2023/11/28 |
| WxB-24 | Tube | 2023/10/10 | 2023/11/14 | . | . | . | 2023/12/26 |
| WxB-25 | Tube | 2023/10/10 | 2023/11/14 | . | . | . | 2023/11/28 |
| WxB-26 | Tube | 2023/10/10 | 2023/11/14 | 2024/04/22 | 2024/05/07 | 2024/05/20 | Breeding |
| WxB-27 | Tube | 2023/10/13 | 2023/11/14 | . | . | . | 2023/12/19 |
| WxB-28 | Tube | 2023/10/13 | 2023/11/14 | . | . | . | 2023/12/19 |
| WxB-29 | Tube | 2023/10/13 | 2023/11/14 | . | . | . | 2023/12/26 |
| WxB-30 | Tube | 2023/10/13 | 2023/11/14 | . | . | . | 2023/12/19 |
| WxB-31 | Tube | 2023/10/20 | 2023/11/14 | . | . | . | 2023/12/19 |
| WxB-32 | Tube | 2023/10/10 | 2023/11/21 | . | . | . | 2024/04/22 |
| WxB-33 | Tube | 2023/10/10 | 2023/11/21 | . | . | . | 2023/12/19 |
| WxB-34 | Tube | 2023/10/10 | 2023/11/21 | . | . | . | 2024/04/22 |
| WxB-35 | Tube | 2023/10/10 | 2023/11/21 | . | 2024/1/4 | . | No data |
| WxB-36 | Tube | 2023/10/10 | 2023/11/21 | . | . | . | 2024/03/26 |
| WxB-37 | Tube | 2023/12/12 | 2024/01/05 | . | . | . | No data |
| WxB-38 | Tube | 2023/12/12 | 2024/01/05 | . | . | . | 2024/02/20 |
| WxB-39 | Tube | 2023/12/12 | 2024/01/12 | 2024/04/22 | 2024/05/07 | 2024/05/21 | Breeding |
| WxB-40 | Tube | 2023/12/12 | 2024/01/12 | 2024/04/22 | 2024/05/10 | 2024/05/24 | Breeding |
| WxB-41 | Tube | 2023/12/12 | 2024/01/16 | 2024/04/22 | 2024/05/07 | 2024/05/20 | Breeding |
| WxB-42 | Tube | 2023/12/12 | 2024/01/19 | 2024/04/22 | 2024/05/07 | 2024/05/20 | 2024/05/21 |
| TwG1-J198 | Combination | 2025/02/14 | 2025/02/28 | 2025/04/01 | 2025/05/28 | No data | Breeding |
| TwG1-J218 | Combination | 2025/02/19 | 2025/03/07 | 2025/04/08 | 2025/04/14 | No data | No data |
| TwG1-J250 | Combination | 2025/02/25 | 2025/03/11 | 2025/04/09 | 2025/04/14 | No data | No data |
| TwG1-J280 | Combination | 2025/03/04 | 2025/03/19 | 2025/04/16 | 2025/04/25 | 2025/05/09 | Breeding |
| TwG1-J283 | Combination | 2025/03/04 | 2025/03/19 | 2025/04/16 | 2025/04/25 | 2025/05/09 | Breeding |
| TwG1-J292 | Combination | 2025/03/04 | 2025/03/19 | 2025/04/16 | 2025/04/28 | No data | Breeding |
| TwG1-J294 | Combination | 2025/03/04 | 2025/03/19 | 2025/04/16 | 2025/04/21 | 2025/05/09 | Breeding |
| TwG1-J295 | Combination | 2025/03/04 | 2025/03/19 | 2025/04/16 | 2025/04/21 | 2025/05/09 | Breeding |
| TwG1-J316 | Combination | 2025/03/07 | 2025/03/21 | 2025/04/17 | 2025/05/02 | No data | Breeding |
| TwG1-J318 | Combination | 2025/03/07 | 2025/03/21 | 2025/04/17 | 2025/04/30 | No data | Breeding |
| TwG1-J321 | Combination | 2025/03/07 | 2025/03/21 | 2025/04/17 | 2025/04/28 | 2025/05/09 | Breeding |
| TwG1-J330 | Combination | 2025/03/11 | 2025/03/25 | 2025/04/22 | 2025/05/07 | No data | Breeding |
| TwG1-J368 | Combination | 2025/03/18 | 2025/04/01 | 2025/04/28 | 2025/05/09 | 2025/06/02 | Breeding |
| TwG1-J333 | Combination | 2025/03/14 | 2025/03/31 | 2025/04/25 | 2025/05/12 | No data | Breeding |
| TwG1-J370 | Combination | 2025/03/21 | 2025/04/04 | 2025/05/01 | 2025/05/07 | 2025/06/02 | Breeding |
| TwG1-J372 | Combination | 2025/03/18 | 2025/04/04 | 2025/05/01 | 2025/05/12 | No data | Breeding |
| TwG1-J373 | Combination | 2025/03/18 | 2025/04/04 | 2025/05/01 | 2025/05/28 | No data | Breeding |
| TwG1-J388 | Combination | 2025/03/25 | 2025/04/04 | 2025/05/08 | 2025/05/21 | No data | Breeding |
